# Cortical potentials evoked by stimulation of cervical vagus vs. auricular nerve: a comparative, parametric study in nonhuman primates

**DOI:** 10.64898/2025.12.30.697054

**Authors:** I. Rembado, M. Ravan, M. Akerman, M.M. Sanchez, K. J. Bascoc, C. Birch, H. Boyd, B. Amoeni, A. Morse, I. Kemp, J. W. Hur, S. Perlmutter, D. Su, C. Sison, E. E. Fetz, S. Zanos

## Abstract

Stimulation of sensory vagal pathways is typically delivered via invasive, cervical vagus nerve stimulation (cVNS) or noninvasive, trans-auricular nerve stimulation (taNS). While both methods are investigated therapeutically, their effects on brain physiology remain poorly understood, hindering mechanistic understanding and stimulus optimization. In 6 awake nonhuman primates, we recorded cortical vagal-evoked potentials (VEPs) from subdural electrodes placed in prefrontal, sensorimotor and parietal cortical areas, in response to cVNS or taNS. Across 478 different taNS and cVNS protocols, we varied stimulation side, intensity, frequency, pulse count, and pulse width (PW) and assessed independent effects on amplitude and latency of early (EC; 30-100 ms), intermediate (IC; 101-200 ms) and late components (LC; 201-500 ms) of VEPs. Fixed and random effects of stimulation parameters and subjects, respectively, on VEP measurements, were assessed using a linear mixed-effects model. Overall, cVNS elicits more robust VEPs than taNS, with larger EC, IC and LC amplitudes in both hemispheres. cVNS-elicited ECs and LCs are largest in PFC and PC areas, whereas ICs are largest in SM areas. On the other hand, taNS generally does not elicit area-specific responses. cVNS-elicited ECs have slower latency than ta-NS elicited ECs. Higher stimulation frequencies and intensities and a longer pulse width elicit larger ECs and ICs for cVNS, and to some extent for taNS. Both short and long cVNS trains elicit stronger ECs, and long trains elicit slower ICs. Earlobe stimulation elicits VEPs that partially overlap with those from taNS. In conclusion, cVNS and taNS elicit cortical VEPs in a manner consistent with distinct engagement of ascending vagal pathways, and with similarities and differences in the effects of stimulation parameters on evoked responses.

## Introduction

The vagus nerve (VN) is a mixed, sensory and motor, mostly autonomic nerve, mediating bidirectional communication between visceral organs and the brain to regulate physiological [1] and immunological homeostasis [2]. The VN is implicated in the pathogenesis of neurological disorders, including epilepsy, depression and recovery from neural injury, and of several chronic non-neurological diseases, including hypertension, heart failure, diabetes, obesity, and certain cancers [3]. For those reasons, vagus nerve stimulation (VNS) is used as treatment in epilepsy [4] and treatment-resistant depression [5], and tested for the treatment of a variety of other diseases, including post-traumatic stress disorder [6] chronic headaches [7], [8], Alzheimer’s disease [9], treatment-resistant anxiety disorders [10]-[12], rehabilitation after stroke [13] or spinal cord injury [14], cardiovascular diseases [15], rheumatoid arthritis [16], inflammatory bowel disease [17] as well as for cognitive augmentation [18]. In many of these clinical applications, VNS targets the sensory vagal pathway, as the ultimate therapeutic target is either a brain circuit or a peripheral organ innervated by an autonomic reflex, that can be activated via VNS through its sensory arm.

Stimulation of the sensory vagal pathway is typically done with electrodes implanted at the cervical level, termed cervical VNS (cVNS). At the cervical level, the VN contains thousands of sensory and motor fibers innervating the heart, lungs, airways, and gastrointestinal tract [19], the majority of which are sensory [19]-[21]. Stimulation of sensory vagal fibers activates first order sensory neurons in the vagal ganglia, e.g. the nodose ganglion, and second order sensory neurons in brainstem nuclei, e.g., the nucleus tractus solitarius (NTS); those neurons then project to other brain sites, including brainstem nuclei, limbic and forebrain sites [22], as well as the thalamus and different cortical areas, where some of the therapeutic effects of VNS are presumed to take place [23]. A less invasive approach to stimulating the sensory vagal pathway is with trans-auricular nerve stimulation (taNS), delivered using noninvasive electrodes. taNS activates a relatively small number of cutaneous afferents innervating part of the skin of the auricle. Because, at least some of, those afferents also terminate on NTS neurons [24], it has been theorized that taNS can in principle activate vagal projections to brain areas that are, at least partially, overlapping with those activated by cVNS, including the cortex. Even though the neuroanatomical and neurodevelopmental similarities and differences between the cervical vagus and its auricular branch are fairly well understood [25], [26], how cVNS and taNS compare with regard to effects on cortical physiology, is unclear. In addition, whether, and how, stimulus parameters of cVNS and taNS affect cortical function is largely unknown. Such knowledge would be relevant to the mechanisms of action of VNS, to VNS parameter optimization and therapy personalization, as well as to gaining insights into the role of the sensory vagal pathway in brain function.

One way to assess effects of VNS on cortical physiology is recording of vagal evoked potentials (VEPs). VEPs are produced in the brain by synchronous ascending volleys elicited on vagal afferents by VNS. The exact neuronal correlates of VEPs are unknown but they likely reflect stimulus-evoked activation of areas in the brainstem, hippocampus, thalamus, and cerebral cortex [24]-[30]. VEP shape, including amplitude and latency of the different components, depends on stimulus parameters like intensity, pulse width, pulsing frequency, as well as brain state at the time of stimulation [31]-[33]. For those reasons, and because of their superior temporal resolution, portability and cost-effectiveness compared to fMRI [34]-[35], VEPs can in principle be used as markers of engagement of vagal neurons and brain circuits, to titrate and personalize VNS or taNS treatments for brain conditions [36]. However, there are no systematic studies documenting the independent effect of VNS parameters on VEPs, elicited invasive or non-invasively via cVNS or taNS, respectively.

Nonhuman primates (NHPs) are useful preclinical models for studying brain responses to sensory nerve stimulation [37]. Recently, studies in NHPs have investigated how cVNS and taNS impact behavioral performance and electrical brain activity [38]. In our study, conducted in awake NHPs, cortical VEPs were recorded, generated by electrical stimulation of the VN, using cVNS, or of the auricular nerve, using taNS, with varying stimulation parameters, including intensity, pulse count, frequency, and pulse width. We found systematic relationships between specific parameters and amplitudes and latencies of VEP components, with implications for the mechanistic understanding of central mechanisms of action of VNS. These parametric relationships were typically different between cVNS and taNS, suggesting that the two stimulation modalities may engage distinct brain circuits in the sensory vagal pathway.

## Materials and Methods

### Subjects

The experiments involved 6 male rhesus macaque monkeys, aged 5 to 8 years, and weighing 8-12.3 kg. The information about the age and weight of each monkey is shown in Table 1. All handling, training, surgical procedures, and housing were approved by the Institutional Animal Care and Use Committee at the University of Washington, and all procedures conformed to the National Institutes of Health Guide for the Care and Use of Laboratory Animals.

**Table 1.**
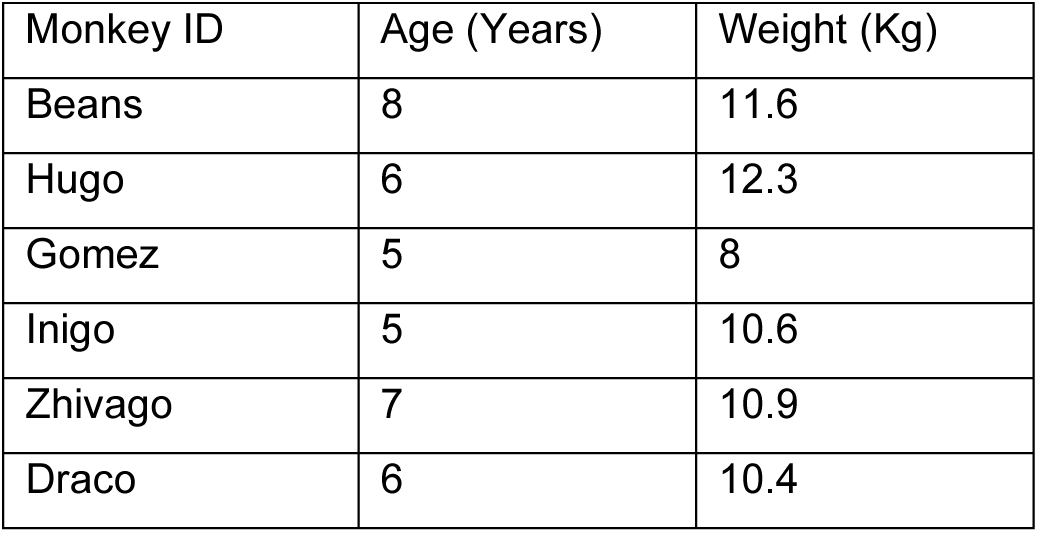
Demographic information of the non-human primates enrolled in the study.

| Monkey ID | Age (Years) | Weight (Kg) |
| --- | --- | --- |
| Beans | 8 | 11.6 |
| Hugo | 6 | 12.3 |
| Gomez | 5 | 8 |
| Inigo | 5 | 10.6 |
| Zhivago | 7 | 10.9 |
| Draco | 6 | 10.4 |

### Cortical Implants

Animals underwent anesthesia using sevoflurane gas. Under sterile conditions, the custom-made dual-electrodes were then surgically implanted. The dual-electrodes were constructed using 5 mm and 3 mm rods, each made of 0.002-inch bare temper hard platinum-iridium wire (Advent Research Materials, Oxford, England) (Fig. 1, A). Each rod was soldered to 34-gauge insulated lead wires, and they were insulated by a 10 µm layer of parylene applied by the University of Washington Microfabrication Facility. The rod tips were de-insulated using a scalpel to reach an impedance between 10 and 50 kΩ (at 1000 Hz). The 3 mm and 5 mm rods were paired in a 0.055-inch polyimide medical (Microlumen, Oldsmar, FL) tubing and secured using silicon sealant (Dow Corning 734, McMaster-Carr) to create a dual-electrode. The tip of the 3 mm rod was placed ∼0.5 mm from the edge of the polyamide tubing, while the tip of the 5 mm rod was placed at 2.0-2.5 mm from the edge. This allows for the 3mm rod to rest on the surface of the brain while the 5 mm rod penetrates the cortex, targeting layer 5. In animals Bean, Gomez, Hugo, and Inigo, the back ends of the lead wires for each dual-electrode were soldered to three printed circuit boards (PCBs) manufactured by Omnetics; there were 16 dual-electrodes attached to each PCB. The three PCBs were then attached to a Blackrock connector (Blackrock Neurotech) inserted in a titanium pedestal through 4 cm wire bundles (Fig. 1, A). A custom-made titanium casing was instead used for Drago and Zhivago animals.

**Figure 1.**
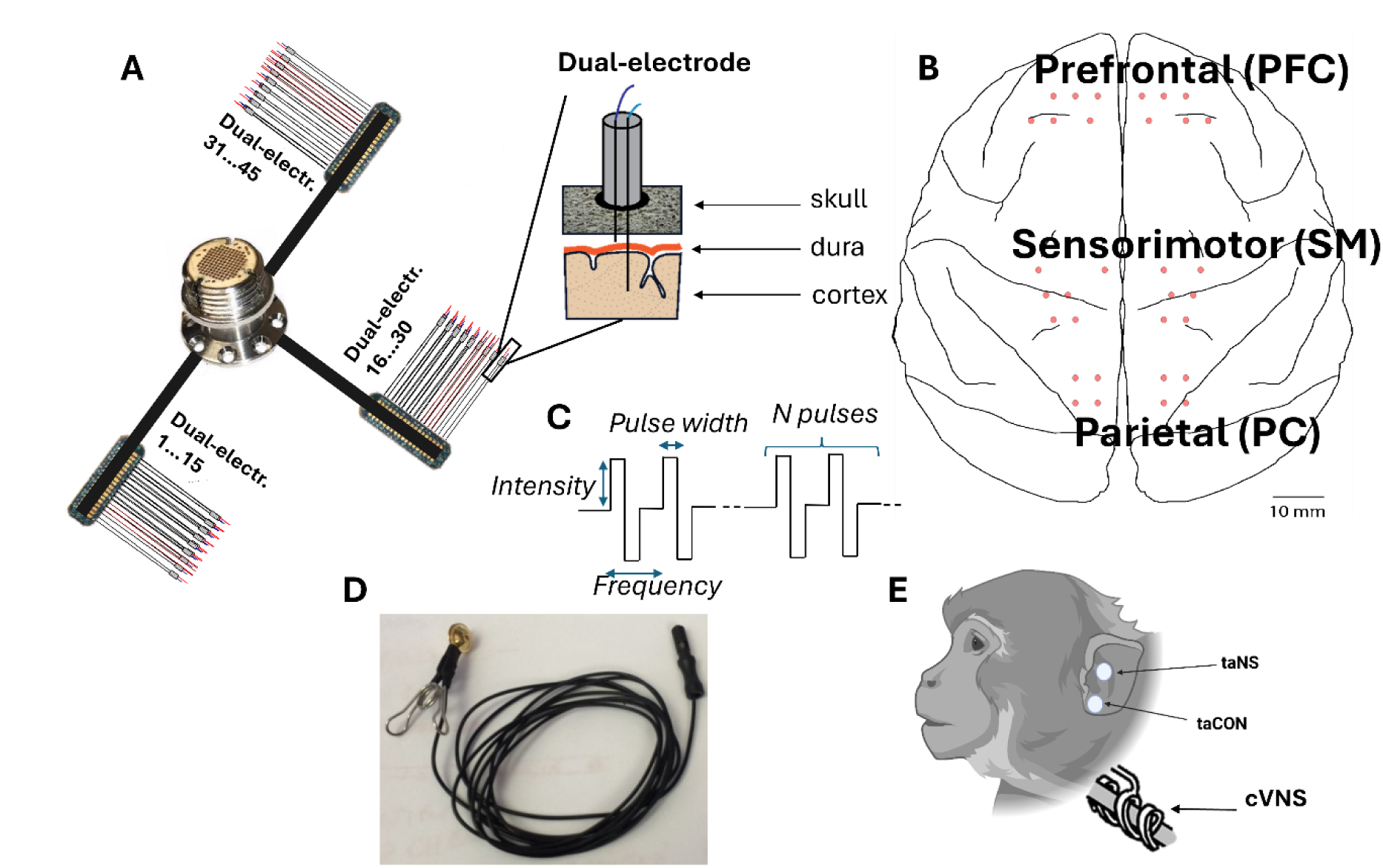
Experimental cortical and vagus nerve implant. (A) Schematic illustration of the dual-electrode implant system with 3 mm and 5 mm rods penetrating the brain. (B) Locations of implanted electrodes on the cortical map. (C) Schematic representation of the electrical stimulation parameters used to generate the protocols investigated in the study. (D) Ear clip used to non-invasively stimulate the auricular branch of the vagus nerve (taNS). (E) Schematic illustration of the location of the stimulated areas for taNS, cVNS and trans-auricular control (taCON). For cVNS the dual helical stimulating cuff was wrapped around the vagus nerve. Created in BioRender. Rembado, I. (2026) https://BioRender.com/b8tkx6h

All animals received postoperative courses of analgesics and antibiotics. The number of implanted electrodes across both hemispheres for each monkey is provided in Table 2. Fig. 1, B provides a schematic illustration of the electrode positions on the cortical map, showing placements over the prefrontal (PFC), sensorimotor (SM), and parietal (PC) cortical areas. The actual number and precise locations of the implanted electrodes for each of the six monkeys are detailed in Fig. S1, illustrating individualized implantation patterns across both hemispheres.

**Table 2.** Cortical electrodes implanted for each animal.

| Monkey | PFC (L/R) | SM(L/R) | PC(L/R) |
| --- | --- | --- | --- |
| Beans | (14/14) | (22/22) | (6/8) |
| Hugo | (16/12) | (16/18) | (12/12) |
| Gomez | (14/14) | (16/18) | (12/12) |
| Inigo | (10/10) | (12/12) | (10/10) |
| Draco | (12/12) | (12/12) | (8/8) |
| Zhivago | (12/12) | (10/10) | (8/8) |

Abbreviations: PFC= prefrontal cortex, SM= sensorimotor, PC= Parietal cortex, L=Left hemisphere, R=Right hemisphere.

#### Monkeys Bean, Hugo, Gomez, Inigo

An incision was made along the midline of the scalp and muscle, and connective tissue were resected to expose the skull. Dual electrodes were placed through 0.5 mm burr holes drilled with a stereotaxic guide. The placement was carried out using forceps to place the electrode in the hole until resistance was felt due to contact of the longer rod with the dura. The longer rod was then pushed through the dura until resistance was felt again, indicating that the longer rod had entered the brain, and the shorter rod was resting on the dura (Fig. 1, A). Between 43 and 32 dual-electrodes (corresponding to 86 and 64 single electrodes) were implanted over PFC, SM, and PC (Fig. 1, B); Figs. S1, A-D). Six skull screws used as ground and reference leads were placed on the occipital or the temporal bone, depending on skull exposure during surgery and the availability of space after the electrode implantation. Acrylic was used to seal the burr holes and hold the electrodes in place. The Blackrock pedestal connector was then screwed with 8 skull screws into the skull. The skin was closed over the lead wires and PCBs, leaving only the Blackrock pedestal containing the connector exposed.

#### Monkeys Draco, and Zhivago

An incision was made along the midline of the scalp and the muscle and connective tissue were resected to expose a sufficient amount of skull for a 2.5-inch diameter titanium casing to be placed. Four to eight screws were placed in the skull around the resected area, at least four of which were T-screws used as grounds and references. Dual electrodes were placed through 0.5 mm burr holes drilled with a stereotaxic guide. 32 electrodes were placed in PFC, SM, and PC areas for animal Draco (Fig. S1, E) and 30 dual electrodes were placed in these areas for animal Zhivago (Fig. S1, F). These animals received EOG electrodes as described above during a second surgery conducted several months after the brain implant surgery. Acrylic was used to seal the burr holes and hold the electrodes in place. The casing was then placed over the implant and secured to the skull screws using acrylic.

### Cervical vagus nerve implants

In a separate procedure, the monkeys underwent the implantation of a stimulating cuff on the cervical VN (Fig. 1, E). Left cervical VN was implanted for animals Draco and Zhivago and the right cervical VN was implanted for Gomez and Inigo. While under general anesthesia with sevoflurane gas, each animal was positioned supine. A horizontal incision was made above the left or right clavicle, medial to the sternocleidomastoid muscle. The skin was retracted to expose the carotid sheath vertically. Subsequently, the sheath was opened, and the vagus nerve was exposed along a length of approximately 3–4 cm between the jugular vein and the carotid artery.

To facilitate manipulation, a rubber loop was threaded behind the nerve, and the VN electrode was positioned around the VN trunk. The proximal end of the electrode leads was then secured with sutures in the subcutaneous tissue near the implant site, providing support for the placement of the nerve cuff on the nerve trunk. The distal end of the leads was routed subcutaneously to a skin opening at the back of the head, near the edge of the head chamber previously installed during the cortical procedure (refer to cortical implant). The leads immediately entered the chamber and were affixed to the base using acrylic. A 2-channel connector facilitated the electrical connection of the cuff leads to a stimulator. Following the procedure, both head and neck incisions were sutured, and the monkey received post-surgery analgesics and antibiotics for a 10-day recovery period.

The bipolar nerve cuff (Cyberonics, LivaNova Inc.) consisted of two platinum–iridium filament bands, each embodied in a flexible, silicone-based, helical loop. A third helical loop was included to stabilize the array around the nerve. The overall length of the cuff, including the three helical loops, measured 3 cm with an internal diameter of 2 mm. After the animal recovered we assembled the small adapter needed to interface with the stimulator and performed the impedance measurements. The impedance between the cuff contacts ranged from 10 kΩ to 30 kΩ for all animals. We also verified that stimulation did not induce neck muscle twitches; when it did, we considered the cuff unusable, as the stimulation was not being delivered selectively to the vagus nerve. For two of the six animals, we were unable to perform cVNS either because the stimulation produced muscle twitches or due to surgical complications or postoperative skin infections along the path of the wires from the neck to the titanium head chamber, forcing us to remove the whole implant.

### Auricular stimulation electrodes

Non-invasive VNS was administered to the auricular branch (taNS) using an ear clip positioned at the cymba conchae (Fig. 1, D, E). Left taNS was tested for Bean, Hugo, Gomez, Draco, and Zhivago, and right taNS was tested for Bean, Hugo, Gomez, and Inigo. As a control for sensory activation, we performed trans-auricular control (taCON) stimulation by delivering stimulation with the same ear clip to the earlobe innervated by the trigeminal nerve (Fig. 1, E). This allowed us to compare the evoked neural responses between VN activation and sensory activation. Left taCON was tested for Hugo, Draco, and Zhivago, and right taCON was tested for Bean. For all sessions to improve electrical conduction, a conductive gel was applied between the two faces of the ear clip.

### Physiology experiments

During daily experimental sessions, each monkey sat in a primate chair inside a soundproof booth, with both forearms comfortably restrained resting at the sides and elbows flexed. The animals were rewarded with fruit sauce for remaining relaxed and avoiding struggling or headshaking given that the head was not restrained. This was particularly important during taNS sessions, as the monkeys required training to tolerate the ear clip. Early training focused on simply keeping the ear clip in place without head shaking; stimulation was then gradually introduced, beginning with low intensities and progressively increasing to higher levels. Over approximately 4–6 weeks of daily training, all animals learned to tolerate the ear clip and stimulation, although occasional head shaking during a session sometimes necessitated aborting the session. To limit the amount of stimulation the monkey received each day, daily sessions were restricted to one hour, allowing us to test up to five protocols per session.

When stimulation was delivered via cVNS, the primary adverse reaction observed was coughing. In such cases, stimulation was immediately halted, and the amplitude and/or frequency was reduced before resuming. These behavioral sensitivities varied across individuals, which partly accounts for differences in stimulation parameters across subjects.

#### Stimulation parameters

taNS, taCon, and cVNS protocols varied in current intensity, pulse count delivered within each train, frequency (Hz, pulse count per train in each second), and the pulse width (Fig, 1 C). The intensities investigated varied from 500 µA to 3250 µA. The pulse count varied from 1 to 60. The frequency of pulses within the train ranged from 0 Hz to 1000 Hz, where 0 Hz is considered for the case where a single pulse was delivered. Trains were delivered at intervals ranging from 0.5 to 10 seconds, with the number of trains varying from a minimum of 4 to a maximum of 318. The pulse widths were either 100 µs or 200 µs. To further analyze the data we organized the stimulation parameters in categories (see statistical methods for details) resulting in 112 protocols recorded from a total of 800 sessions. The total number of sessions and protocols, along with the corresponding parameters for each monkey, are summarized in Table 3 and reported extensively in supplementary Table S1. Stimulation was delivered using a STG4008 stimulator (Multichannel Systems).

**Table 3.** Stimulation protocols in different animals.

| Animal | N. of Sessions | N. of protocols | Stimulation modalities | Intensity (μA) | Pulse count | Frequency (Hz) | Pulse width (μs) |
| --- | --- | --- | --- | --- | --- | --- | --- |
| Beans | 125 | 25 | Left taNS,<br>Right taNS,<br>Right taCON | 500,1000, 1250,<br>1500, 2000, 2500,<br>3000 | 1, 4, 10, 15, 20,<br>60 | 0, 30, 50,<br>100, 200, 300 | 100,<br>200 |
| Hugo | 130 | 18 | Left taNS,<br>Right taNS,<br>Left taCON | 500,1000, 1200,<br>1250, 1500, 2000,<br>2500, 3000 | 1, 4, 10, 15, 20,<br>60 | 0, 30, 50,<br>100, 200, 300 | 100,<br>200 |
| Gomez | 150 | 27 | Left taNS,<br>Right taNS,<br>Right cVNS | 500, 800, 1000,<br>1250, 1400, 1500,<br>1600, 2500, 3000 | 1, 4, 10, 15, 20,<br>60 | 0, 30, 50,<br>100, 200, 300 | 100,<br>200 |
| Inigo | 151 | 21 | Right taNS,<br>Right cVNS | 500, 800,<br>1000,1200,1250,<br>1500,1800 | 1, 4, 10, 15, 20,<br>60 | 0, 30, 50,<br>200, 300 | 100,<br>200 |
| Draco | 154 | 11 | Left taNS,<br>Left taCON,<br>Left cVNS | 500, 750, 800,<br>1000, 1250, 1500,<br>2000 | 4, 5, 7, 9, 10,<br>15 | 50, 200, 300 | 200 |
| Zhivago | 90 | 10 | Left taNS,<br>Left taCON,<br>Left cVNS | 1000, 1250, 1500,<br>1750, 2000, 2250,<br>2500, 2750, 3000,<br>3250 | 4, 5, 7, 9, 10 | 50, 200,<br>300,1000 | 200 |

#### Cortical Recordings

Signals from all cortical electrodes were recorded relative to reference and ground using two 16-channel, DC-coupled g.USB amplifiers (g.tec Medical Engineering Gmbh, Schiedlberg, Austria) for animals Draco and Zhivago. Signals were sampled at 24-bit resolution, at 4.8 Ksamples/sec or 9.6 Ksamples/sec with no filtering. Data from the amplifiers was streamed to a personal computer through a USB link, then stored and visualized in real-time using a Simulink-based (MathWorks, Natick, MA) graphical user interface, developed in-house. Signals from the other animals were recorded using six 16-channel ZC-16 headstages (Tucker-Davis Technologies, TDT, Alachua, FL), routed to a TDT-RZ2 BioAmp Processor and TDT-PZ5 NeuroDigitizer Amplifier (24-bit resolution, sampling rates of 3051.76 samples/sec or 6103.52 samples/sec).

#### Vagal Evoked Potentials (VEPs)

Raw signals from each cortical site were visually inspected, and bad channels were removed. A third-order antialiasing filter was then applied to each raw signal (frequency cut-off Hz: [1 - sampling rate/2-1). The signals were then segmented into two seconds long windows centered around the stimulation onsets from -1 s to 1 s. The average across trials belonging to the same stimulation protocol was then computed, generating the so-called VEPs, considered to be the brain responses to vagus nerve stimulation (VNS). The compiled VEPs for each channel and each stimulation protocol then underwent through an artifact removal and signal denoise pipeline (see details below).

#### Signal preprocessing

We implemented a preprocessing and denoising pipeline applied to all VEPs. The code was custom made in Matlab. The preprocessing pipeline consisted of three major steps:

#### I. Signal quality control (QC)

Signals first underwent electrode-level assessment to remove non-physiological channels based on baseline variability metrics. Channels belonging to the same area were grouped together. For each channel the baseline window was defined from -250 ms to -20 ms from stimulation onset. For each channel, we computed (i) a robust deviation score, based on the mean absolute deviation of that channel’s baseline from the across-channels median baseline, normalized by the standard deviation of the across-channels median baseline, and (ii) baseline variance. Channels were excluded if they showed poor physiology by any of the following criteria: 1) their deviation score exceeded a z-threshold of 2, 2) their variance was <0.2× or >5× the median baseline variance, or 3) they exhibited near-flat activity (variance < 1×10⁻⁶). These thresholds ensured that only physiologically meaningful channels were retained for subsequent artifact-removal procedures. Low-variance channels typically reflect disconnected or poorly contacting electrodes that are unable to capture brain activity, whereas high-variance channels are often contaminated by muscle contractions, amplifier saturation, or environmental noise rather than neural responses. Similarly, large deviations from the median baseline suggest unstable or channel-specific artifacts. Removing such channels prevents unreliable signals from influencing subsequent steps, improving the accuracy and reproducibility of all downstream analyses.

##### II. Stimulation artifact suppression and denoising

Stimulation-related artifacts were removed exclusively within a restricted artifact-processing window spanning from just 20 ms before stimulation onset to 40 ms after the stimulation burst duration. For each channel, candidate artifact spikes were first detected by examining both amplitude and slope excursions relative to pre-stimulus baseline noise. Specifically, samples exceeding an amplitude threshold (5 × baseline standard deviation (SD)) or a derivative threshold based on a robust scale estimate (4 × median absolute deviation (MAD) of first-order differences, scaled by 0.6745 to convert MAD to a standard-deviation–equivalent measure) were marked as putative stimulation pulses. Closely spaced detections were then merged to ensure a single detection per pulse. Around each detected pulse, a short waveform segment from 3 ms before the pulse to 3 ms after the pulse was extracted, and a median artifact template was generated to obtain aa typical stimulation-induced transient. This template was scaled to each occurrence using a bounded least-squares gain (0 ≤ a ≤ 0.8) to prevent over-subtraction and to preserve neural structure. In the response window (0–500 ms), the gain was further down-weighted to 30% to avoid attenuating legitimate cortical activity. The scaled template was then subtracted and the corrected region locally smoothed using a Savitzky–Golay filter (third-order, 11-sample frame) to restore signal continuity.

Following template subtraction, a comprehensive artifact-interpolation step was applied to recover the underlying VEP morphology within the stimulation artifact window. Residual artifact-contaminated samples were identified where the signal exceeded 4× the baseline standard deviation, indicating persistent stimulation energy or amplifier distortion. These samples were grouped into contiguous clusters representing stimulation spike bursts.

For each burst, a ±20-sample neighborhood of the surrounding clean signal was extracted to model local neural dynamics. All corrupt samples were temporarily excluded, and if ≥5 valid samples remained, providing sufficient information to capture local curvature, the missing region was reconstructed using shape-preserving cubic Hermite interpolation (PCHIP). PCHIP was chosen because it maintains the monotonicity and sharp gradients characteristic of neural waveforms while preventing the overshoot and ringing artifacts commonly introduced by standard cubic splines or polynomial interpolation.

To ensure physiologically plausible reconstruction, interpolated values were constrained to the min–max range of neighboring clean data. A short Savitzky–Golay smoothing filter (≤3rd order, ≤11-sample frame) was then applied only within the modified segment to improve continuity without blurring VEP features. Together, these steps enabled accurate recovery of neural responses while preventing artifact-driven distortions.

Next, a secondary hybrid de-spiking step targeted subtle residual spikes not fully captured in the first pass. A robust noise estimate was computed using 1.4826×MAD of the residual signal (a Gaussian-consistent SD estimator). Samples exceeding 3× this robust deviation, grouped with a 1-ms temporal dilation, were marked as artifacts. Short bursts (≤3 samples) were replaced using the local median (ideal for impulsive noise), whereas longer bursts were reconstructed using linear interpolation between the nearest valid samples to maintain continuity. This correction was iterated up to 5 times, or until convergence, to ensure maximum suppression without damaging the neural structure.

To protect authentic neural features at the artifact boundaries, the cleaned signal was blended only at the edges of the artifact window using a 6-ms Hann taper, enabling a smooth, symmetric transition between interpolated and original data and preventing abrupt discontinuities.

Finally, any remaining non-finite values (NaNs) were corrected through a hierarchical imputation process: (1) extending the nearest finite edge value into missing regions; 2) linear interpolation where possible; (3) local median replacement for any isolated leftover points.

##### III. Signal Refinement

A final Savitzky–Golay low-pass smoothing (order 3, frame 101) was applied across the entire channel to suppress high-frequency noise while maintaining the shape of the evoked potentials. Cleaned channels belonging to the same cortical area (PFC, PC, SM) for each hemisphere (left, right) were then averaged to produce a high-fidelity denoised VEP waveform for downstream peak/trough analysis. A final visual inspection of the obtained smoothed and averaged VEPs for each area was then performed and obviously non-physiological signals were further removed from subsequent analyses.

For illustrative purposes, Fig. 2, A shows the raw single channel signals (color traces) recorded from SM cortical area of the left hemisphere when 10 taNS pulses were delivered to the left ear at 1000 µA current intensity, 300 Hz and with a pulse width of 200 µs in animal Bean. After channel quality control, stimulation artifact removal and signal refinement, the signal across channels was averaged to obtain a single signal representing the VEP for that area (black thick trace). From this VEP we detected 3 waves and extracted several features considered for the subsequent statistical analysis.

**Figure 2.**
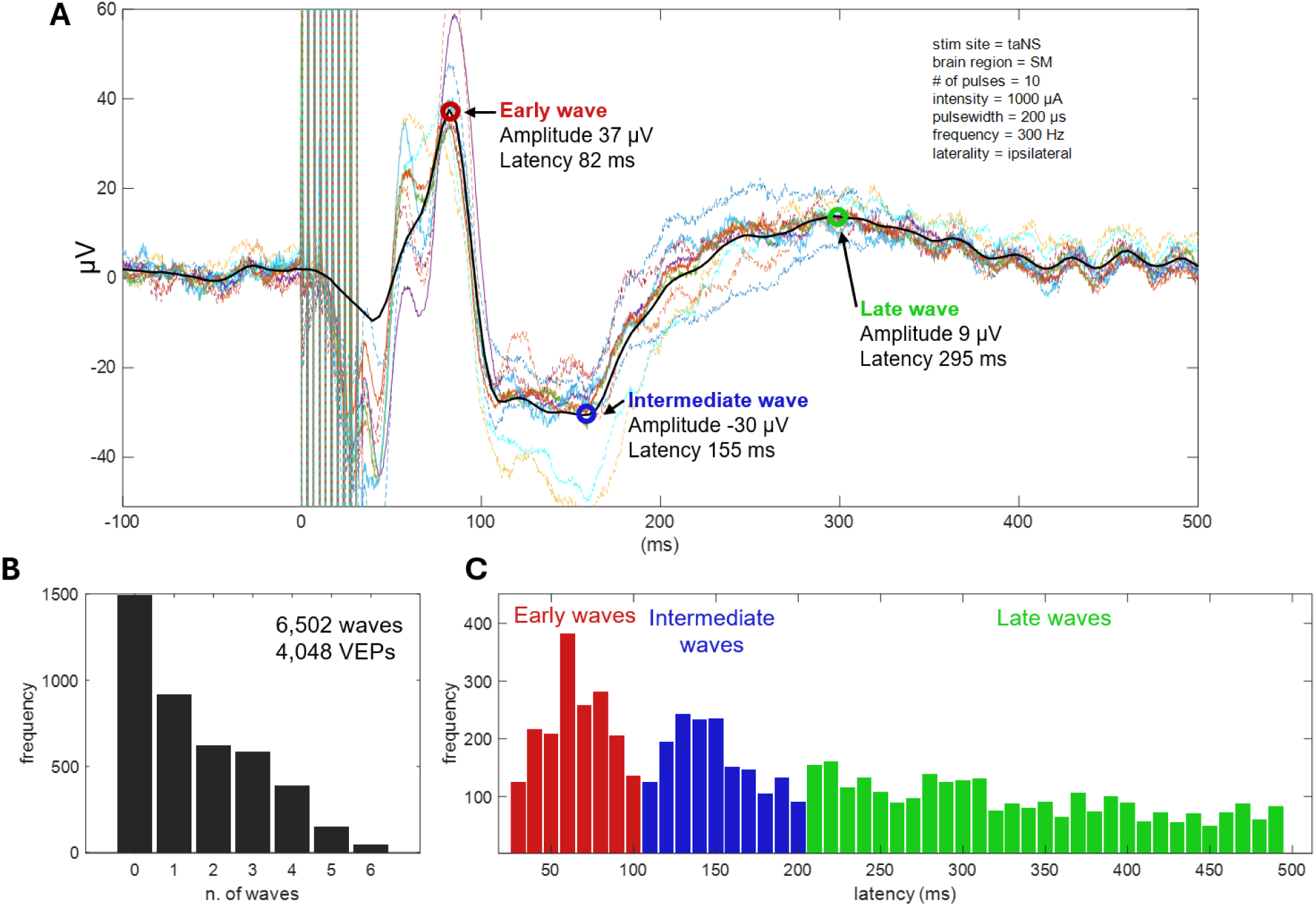
Processing and characterization of vagal evoked potentials (VEPs). (A) Shown are stimulus-evoked potentials across all recording sites in left SM area (colored lines); by averaging across the sites, the SM area-specific vagal-evoked potential (VEP) is calculated, followed by smoothing (black trace). Several features are extracted from the VEP, specifically, the latency and amplitude of waves with amplitudes exceeding three standard deviations above the mean of pre-stimulus baseline. In this example, 3 waves are detected, at 3 different latencies (early, intermediate and late). Stimulation parameters used in this example are shown in the top-right. Note that the absolute value of the amplitude of the detected waves was used in statistical analyses. (B) Distribution of number of waves detected in single VEPs, across all animals and stimulation protocols (6,502 waves from 4,048 VEPs). (C) Distribution of latencies of the detected waves. Based on this distribution, we defined waves occurring 30-100 ms post-stimulus as Early Component (EC), 101-200 ms as Intermediate Component (IC) and 201-500 ms as Late Component (LC). The overall shape of the latency distribution was similar in subsets of VEPs with 1, 2, 3 etc. waves (Suppl. Figure S2).

#### VEP features

To statistically compare cortical potentials evoked from the different stimulation protocols across all six animals, we extracted latency and (absolute) amplitude of waves in the VEPs. We first detected all waves (peaks and troughs) in the VEP trace, within a time window of 30 to 500 milliseconds following stimulus onset. A positive or a negative deflection was considered a valid wave if its absolute amplitude exceeded three times the standard deviation of the pre-stimulus baseline, defined as the interval from 250 ms to 20 ms before stimulation onset.

Detected waves were then categorized into one of 3 VEP components, based on their latency. The choice of latency windows is based on the distributions of latencies of waves detected in the VEPs (Fig. 2B, C; Suppl. Fig. S2), and is in line with our previous study [33].

The 3 components include:

1. an Early Component (EC), occurring between 30 ms and 100 ms after stimulation onset,
2. an Intermediate Component (IC), occurring between 100 ms and 200 ms, and
3. a Late Component (LC), occurring after 200 ms from stimulation onset.

In cases when 2 or more waves were detected within a given latency window, only the largest wave was considered for the statistical analysis. The latency and absolute amplitude of the identified EC, IC, and LC were registered in all VEPs and were then statistically compared across different stimulation protocols and animals.

### Statistical analysis

Following signal processing, we obtained a total number of 4160 VEPs: 2114 generated by taNS, 1934 by cVNS and 112 by taCON (Table S1). Because taCON had a significantly smaller number of trials, we excluded this modality for the overall statistical analyses, although comparisons to taNS are shown (Fig. S3). For each VEP feature (EC amplitude, EC latency, IC amplitude, IC latency, LC amplitude, LC latency), we fitted a linear mixed-effects model (LMM) using restricted maximum likelihood (REML). That approach was followed because multiple recording sessions and stimulation protocols were performed in each animal, with unbalanced representation of protocols across animals, and sparse coverage of specific stimulation parameters within individuals (i.e., an insufficient number of trials for certain numeric values).

Categorical independent variables were defined as

- Stimulation modality: taNS, cVNS (and taCON in a subset of the experiments)
- Stimulation intensity (µA): <1000, [1000–1499], [1500–19999], ≥2000
- Pulse width: 100, 200 µs
- Number of pulses in stimulus train: 1–4, 5–20, >20
- Pulsing frequency (Hz): [0–29], [30–100], >100
- Laterality (stimulation side relative to hemisphere of recorded VEP): ipsilateral, contralateral
- Cortical area of recorded VEP: PC, PFC, SM

In the model, these categorical variables were considered as fixed effects, while the monkey ID as a random effect, to account for within-subject correlation; we also considered two-way interactions between stimulation modality and each fixed effect. Standard assumptions of Gaussian residuals and variance homogeneity were tested, and log transformations were explored, if needed. Least squares means (adjusted means) for each factor level were estimated and compared with Tukey-adjusted pairwise tests. All statistical models were evaluated for convergence using standard diagnostics. Optimization criteria were examined, stability of parameter estimates across iterations was confirmed, and the variance-covariance matrices were verified to be positive definite. No convergence issues were detected, and all models met predefined validity checks. Consequently, all protocols (stimulation modality of taNS or cVNS) were retained in the final analyses.

The intraclass correlation coefficient (ICC) was calculated from variance components estimated in each LMM, as the ratio of animal-level variance to the total variance (animal-level plus residual variance). ICC was used to quantify the proportion of variance attributable to animal-level factors, where values near 1 indicate most variance is due to animal effects and values near 0 indicate minimal animal-level contribution. We also estimated and reported as supplementary material the beta estimates which allowed us to understand how the model is parameterized and how interactions influence the effect of predictors.

A significance threshold of p < 0.05 was used to detect statistical significance. Analyses were performed using SAS version 9.4 (SAS Institute Inc., Cary, NC).

## Results

### Cortical vagal-evoked potentials elicited by taNS vs. cVNS

In 6 animals, the cervical vagus nerve was stimulated invasively via an implanted cuff (cVNS), whereas the auricular nerve was stimulated noninvasively via an ear clip positioned at the cymba conchae (taNS) (Fig.1, D, E). Several cVNS and taNS stimulation protocols were tested, consisting of stimulus trains of different combinations of stimulus parameters, such as intensity, pulse width, pulsing frequency, etc. (Table 3). Using cortical electrodes implanted in both hemispheres, including areas in the prefrontal (PFC), sensorimotor (SM), and parietal cortex (PC) (Fig. 1, A, B), we recorded stimulus-elicited, vagal-evoked potentials (VEPs) (Fig. 2, A). Each VEP was quantified by the amplitude and latency of waves (Fig. 2, A), grouped into 3 components according to their latency: an Early Component (EC; waves with 30-100 ms latency), an Intermediate Component (IC; 101-200 ms latency), and a Late Component (LC; 201-500 ms latency) (Fig. 2, C).

We analyzed a total of 1934 cVNS-elicited VEPs, in 4 animals, and 2114 taNS-elicited VEPs, in 6 animals. Around 35% of VEPs have no identifiable waves, with the remaining having between 1 and 6 waves (Fig. 2, B). After accounting for fixed and random effects of several independent variables, we found that cVNS elicits VEPs with larger amplitude in all 3 components, and slower latency in the EC, compared to taNS (Fig. 3; Table 4). In more detail, the amplitude of the EC is larger with cVNS compared to taNS (Fig. 3, A) and the latency of the EC is longer with cVNS (Fig. 3, B). The amplitude of the IC is larger with cVNS than taNS (Fig. 3, C), even though the IC latency is no different between cVNS and taNS. Finally, the amplitude of the LC is larger with cVNS than taNS (Fig. 3, D), with no difference in the latency between the 2 stimulation modalities. For every comparison, ICC values are relatively small, reflecting a small degree of clustering by animal, indicating that individual differences among animals do not account for a meaningful portion of the observed variance in the VEP features. The beta estimates confirm the statistical significance of these results, providing more insight into the model structure (Suppl. Tables S2, S3, S4, S5, S6, S7).

**Figure 3.**
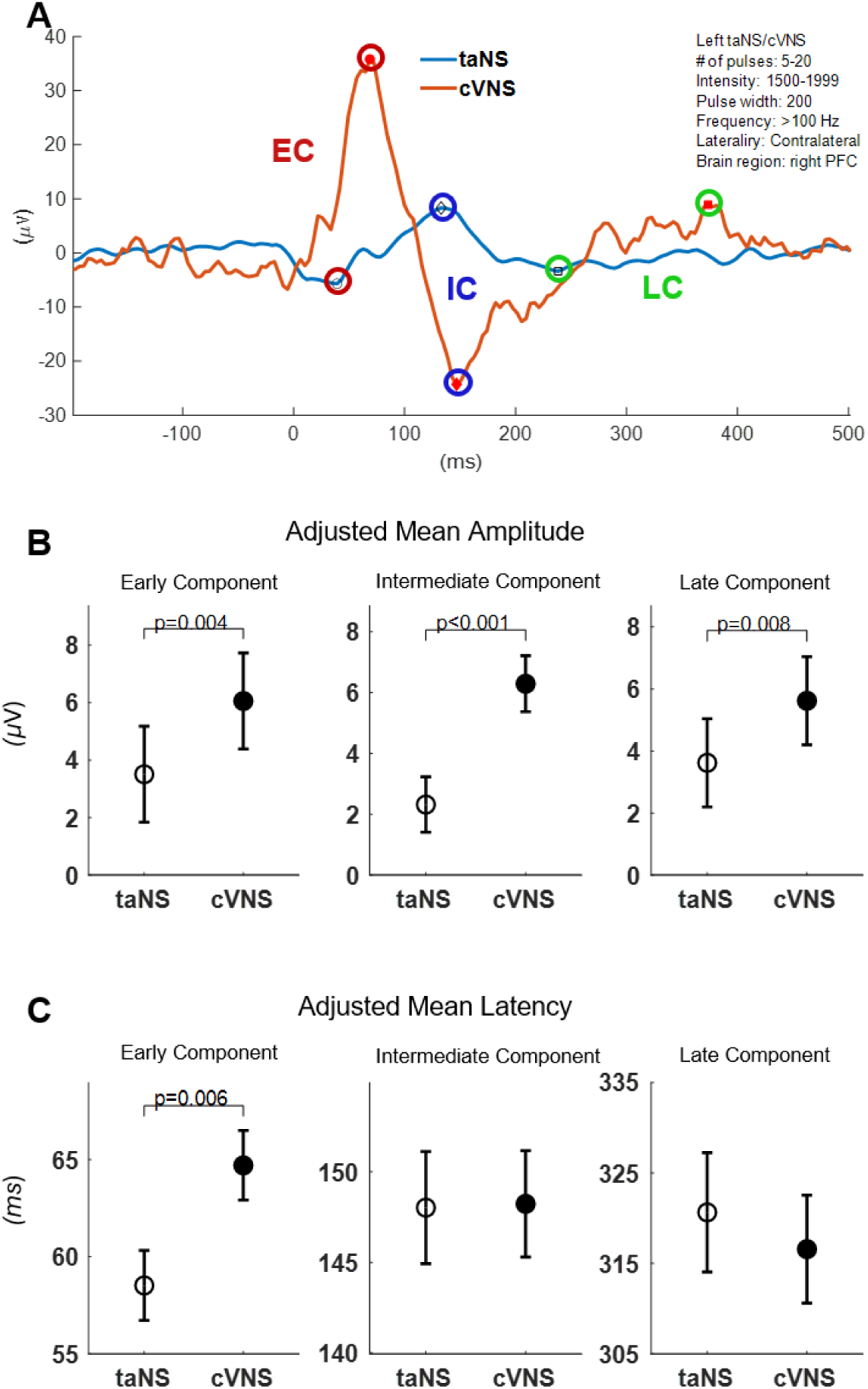
VEPs elicited by cVNS vs. taNS. (Α) Example VEPs elicited by taNS (blue trace) and by cVNS (red trace), of similar stimulation parameters (shown in the top-right); marked are detected early, intermediate and late components (EC, red; IC, blue; LC, green symbols, respectively). (B) Adjusted means (± SEM) for EC, IC and LC amplitude, across all animals and stimulation parameters, for taNS vs. cVNS. (C) Adjusted means for EC, IC and LC latency, across all animals and stimulation parameters, for taNS vs. cVNS.

**Table 4.** Output of the Restricted Maximum Likelihood Linear Mixed-Effects Model.

| Outcome | Factor Level | LS-Mean* | 95% Confidence Interval | p-value | ICC |
| --- | --- | --- | --- | --- | --- |
| EC, Latency (ms) | cVNS | 64.7 | [61.19, 68.21] | 0.0057 | 0.015 |
|  | taNS | 58.52 | [54.98, 62.06] |  |  |
| EC, Amplitude (μV) | cVNS | 6.06 | [2.78, 9.34] | 0.0039 | 0.089 |
|  | taNS | 3.51 | [0.24, 6.79] |  |  |
| IC, Latency (ms) | cVNS | 148.23 | [142.46, 154] | <0.0001 | 0.022 |
|  | taNS | 148.02 | [141.93, 154.11] |  |  |
| IC, Amplitude (μV) | cVNS | 6.29 | [4.49, 8.1] | 0.623 | 0.006 |
|  | taNS | 2.32 | [0.53, 4.1] |  |  |
| LC, Latency (ms) | cVNS | 316.57 | [304.85, 328.29] | 0.9416 | 0.049 |
|  | taNS | 320.64 | [307.69, 333.59] |  |  |
| LC, Amplitude (μV) | cVNS | 5.62 | [2.83, 8.41] | 0.0079 | 0.088 |
|  | taNS | 3.62 | [0.84, 6.4] |  |  |

\* Least square means estimates were adjusted for the random effect of monkey and fixed effects of stimulation modality, brain region, number of pulses, intensity, pulse width, frequency, and laterality.

In a small number of experiments, we compared taNS-elicited VEPs with a common cutaneous control stimulation site in the earlobe (taCON). We found that taCON elicits VEPs with larger EC, IC and LC, and significantly faster IC, compared to taNS (Suppl. Fig. S4). For the amplitude comparisons, ICC values were large, indicating individual differences among animals account for a significant portion of the observed variance; for the latency comparisons, ICC values were small (Suppl. Table S8).

### Parametric effects of cVNS vs. taNS on the early component of cortical VEPs

The Early Component (EC) of VEPs elicited by cVNS is larger than that elicited by taNS (Fig. 3, A-C), with several factors modulating that effect (Fig. 4). EC amplitude for cVNS is greater than taNS for PC and PFC recording sites; in addition, cVNS-elicited PC and PFC ECs are larger compared to SM (Fig. 4, A), whereas for taNS there is no difference across areas (Fig. 4, A). The EC of VEPs is larger for higher frequencies, for both cVNS and taNS (Fig. 4B). Ipsilateral cVNS evokes larger EC responses than contralateral stimulation, in contrast to taNS in which laterality has no effect (Fig. 4, D). EC responses to cVNS exhibit an inverted-U-like relationship with pulse counts (Fig 4, E). Finally, EC responses to both cVNS and taNS increase monotonically with current intensity (Fig 4, F).

**Figure 4.**
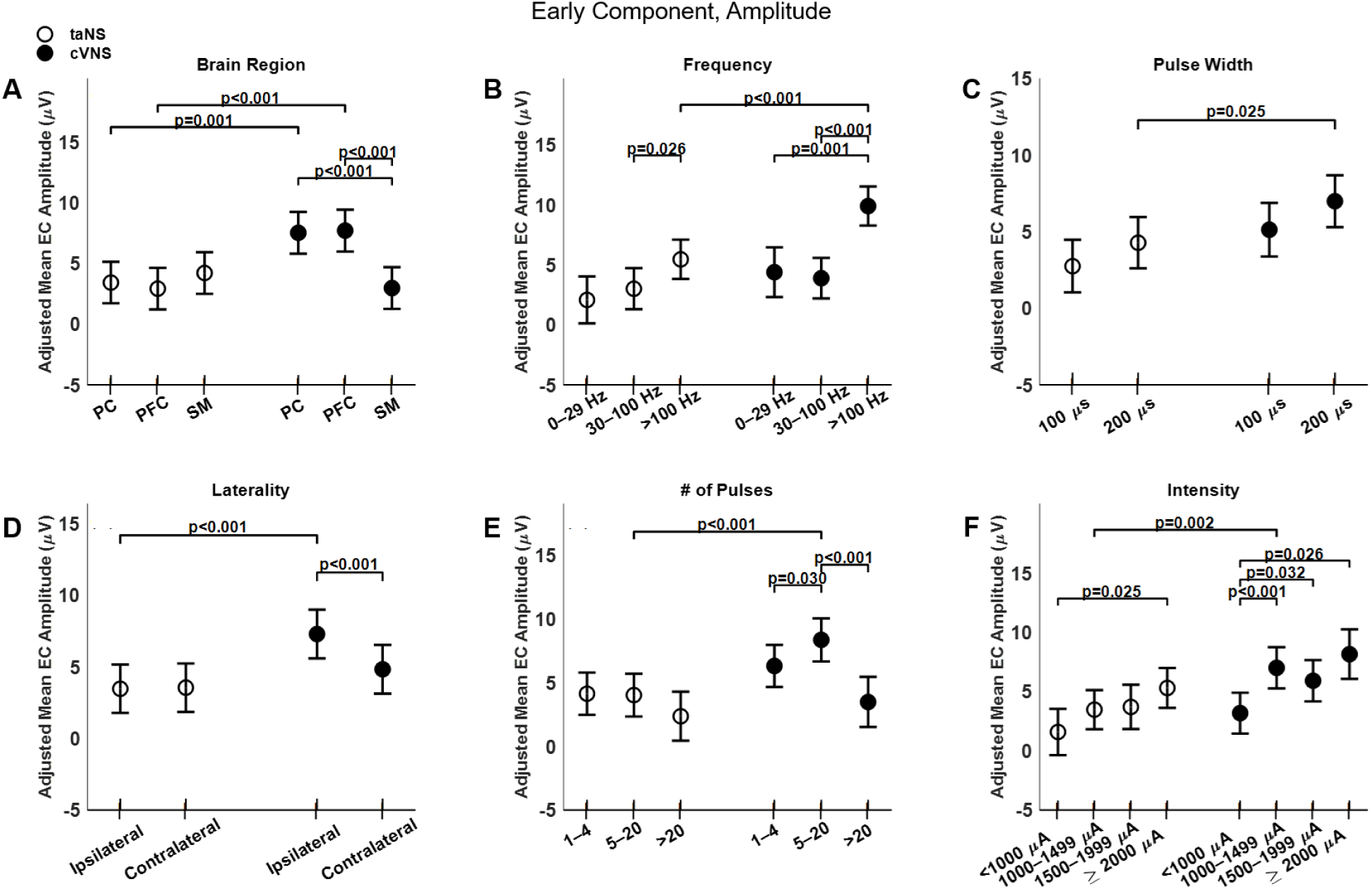
Effects of stimulation parameters on amplitude of the early component (EC) of VEPs. (A) Adjusted mean values (± SEM) of EC amplitude for taNS and cVNS at different brain areas. (B) Adjusted mean values for taNS and cVNS for different stimulation frequencies. (C) Adjusted mean values for taNS and cVNS for different pulse widths. (D) Adjusted mean values for taNS and cVNS for ipsilateral and contralateral stimulation. (E) Adjusted mean values for taNS and cVNS for different pulse counts. (F) Adjusted mean values for taNS and cVNS for different current intensities. In all the plots taNS is shown with empty circles and cVNS with filled circles.

Trans-auricular NS elicits faster ECs compared to cVNS (Fig. 3, D-F), primarily in PFC areas (Fig. 5, A). Higher pulsing frequencies (Fig. 5, B) and a longer pulse width (Fig. 5, C) tend to elicit slower ECs for cVNS, but not for taNS. EC responses to ipsilateral taNS are significantly slower than contralateral taNS (Fig 5, D). Longer cVNS and taNS trains show a, non-significant, trend towards faster EC responses (Fig. 5, E), and intensity does not significantly impact EC latency (Fig. 5, F).

**Figure 5.**
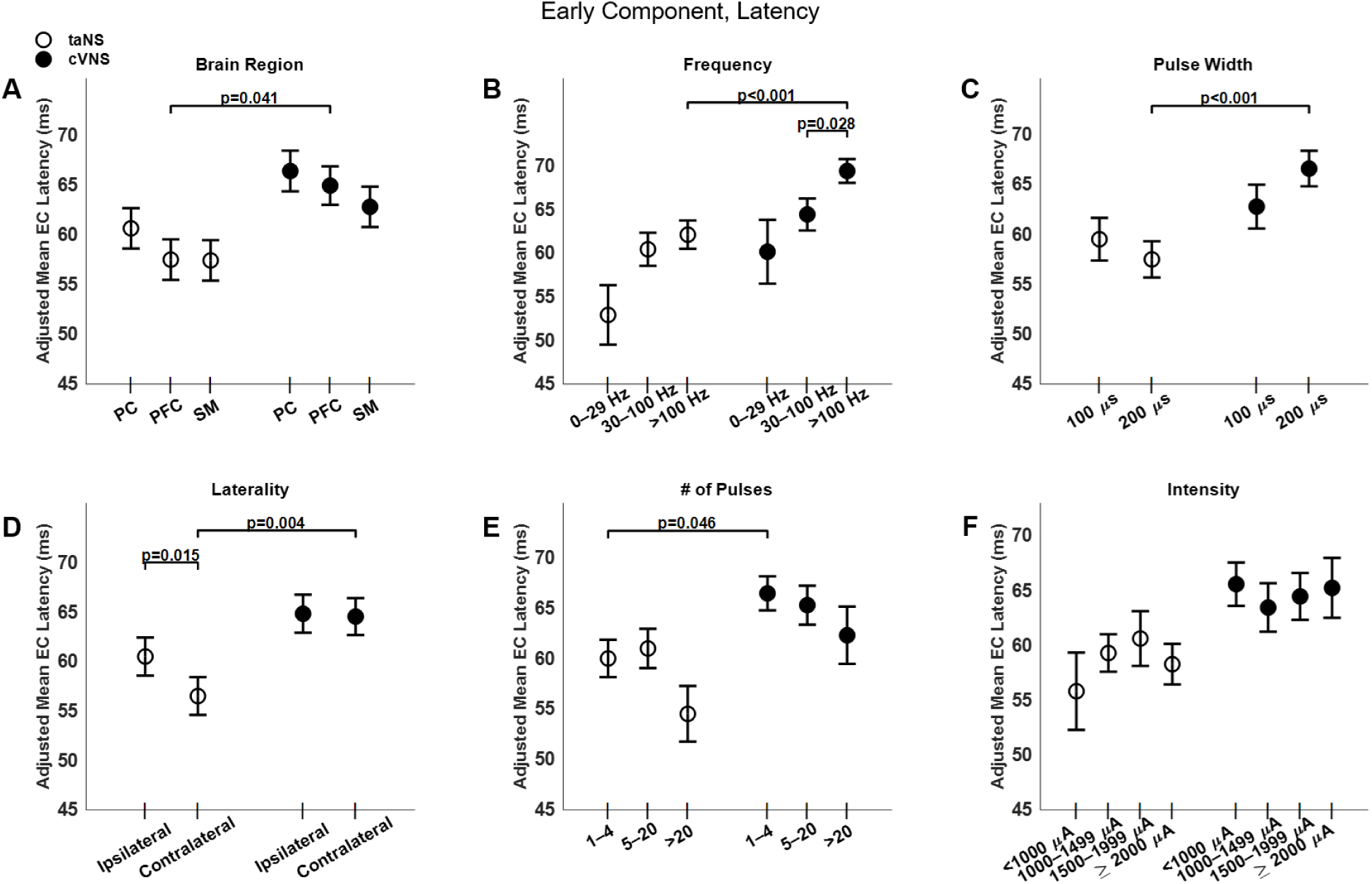
Effects of stimulation parameters on latency of the early component (EC) of VEPs. (A)-(F) Same as Fig. 4, but for EC latency.

All beta estimates for EC amplitude and latency are reported in Tables S2, S3.

### Parametric effects of cVNS vs. taNS on the intermediate component of cortical VEPs

There are notable similarities and differences between cVNS and taNS, with regard to the intermediate component (IC) of VEPs. Overall, IC amplitude for cVNS is larger compared to taNS (Fig. 3, B) across all cortical areas (Fig. 6, A). For cVNS the largest IC responses are in PC, while for taNS the largest responses are over SM area (Fig. 6, A). Stimulation frequency exerts a strong influence on IC amplitude: both cVNS and taNS amplitudes increase with higher frequencies (Fig. 6, B). Both taNS and cVNS responses increase with pulse width (Fig. 6, C). Laterality does not alter the amplitude of the IC, although cVNS is significantly larger than taNS for both ipsilateral and contralateral configurations (Fig. 6, D). Current intensity strongly and positively affects both cVNS- and taNS-elicited IC amplitude (Fig. 6, F).

**Figure 6.**
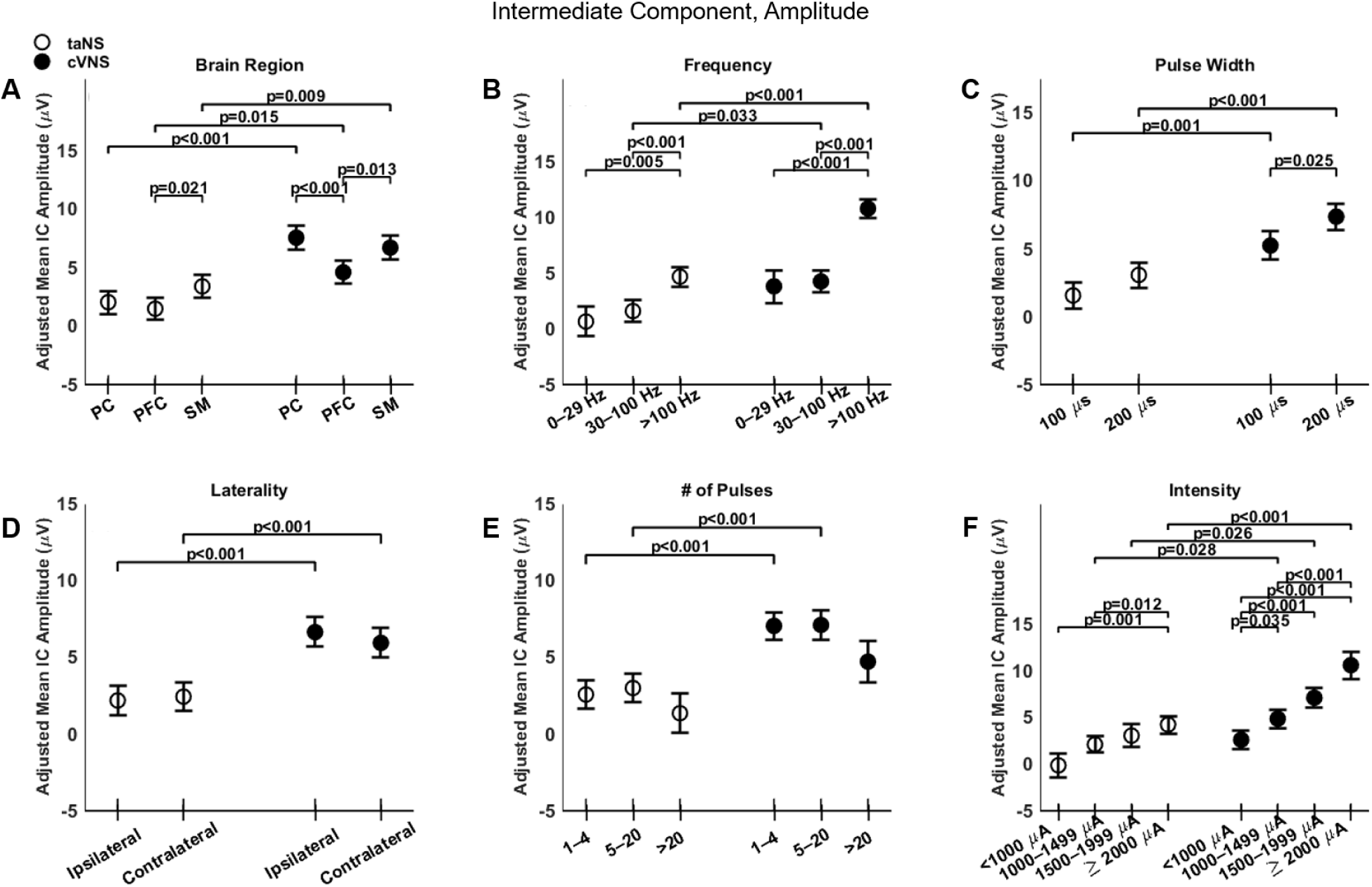
Effects of stimulation parameters on the amplitude of the intermediate component (IC) of VEPs. (A-F) Same as Fig. 4, but for the amplitude of the IC.

The latency of the intermediate component does not show differences between cVNS and taNS (Fig. 3) and few factors affected it (Fig. 7). IC responses in SM area are faster than those in PFC, but only for taNS (Fig. 7, A). Stimulation frequency shows an inverted U-shaped relationship with IC latency with cVNS, and a similar, but non-significant, trend with taNS (Fig. 7, B). Longer cVNS trains are associated with slower IC responses (Fig. 7, C).

**Figure 7.**
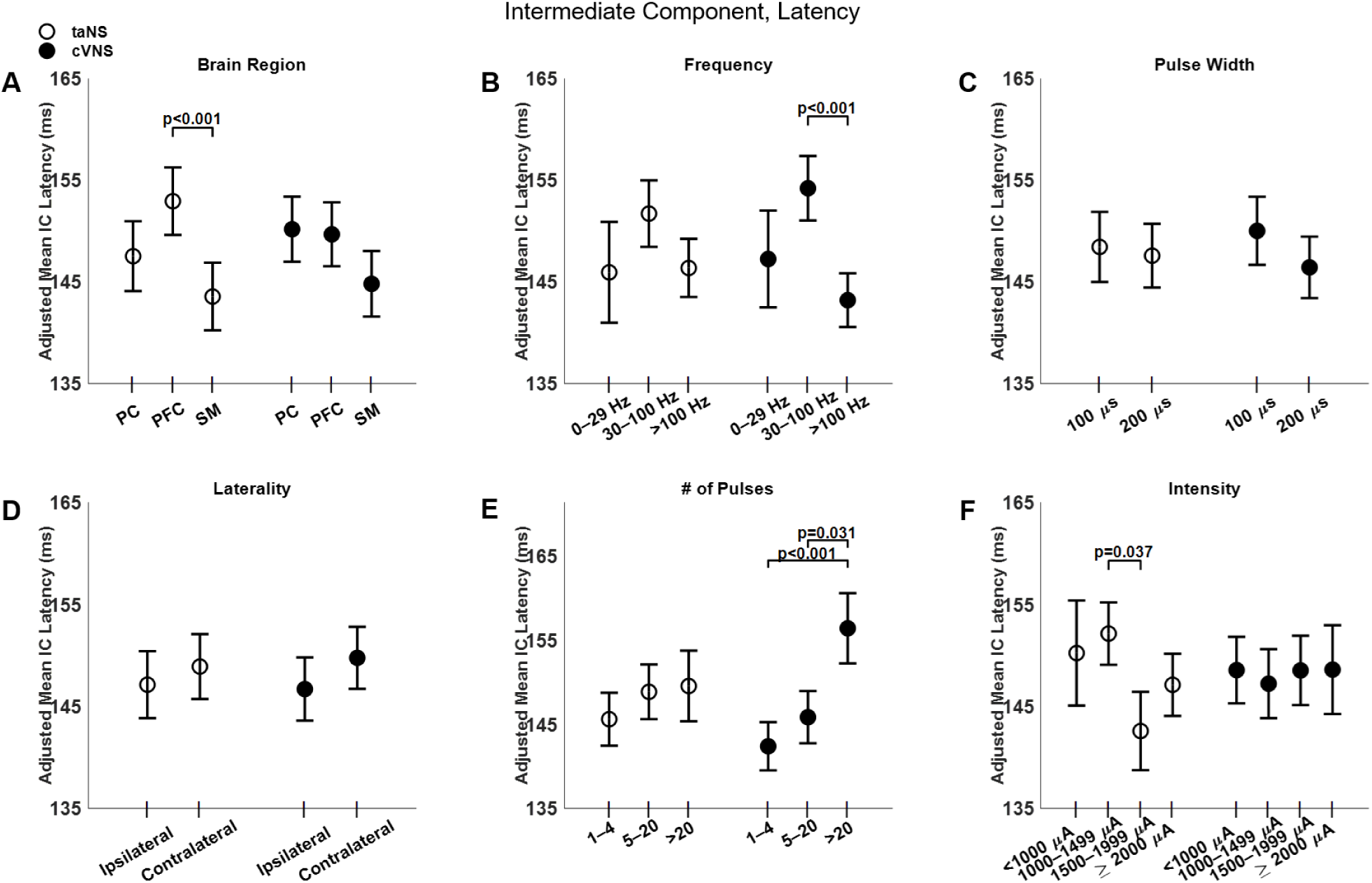
Effects of stimulation parameters on the latency of the intermediate component (IC) of VEPs. (A-F): Same as Fig. 4, but for the latency of the IC.

The beta estimates for the EC amplitude and latency are reported in Tables S4, S5.

### Parametric effects of cVNS vs. taNS on the late component of cortical VEPs

Regarding the late component (LC), cVNS shows consistently larger responses than taNS (Fig. 3). The cortical distributions of the LC follow almost opposite trends between cVNS and taNS (Fig. 8, A). For both taNS and cVNS, IC responses increase with increasing pulsing frequency (Fig. 8, B) and longer pulse width (Fig. 8, C). Laterality and pulse count do not modulate IC responses for neither cVNS nor taNS, although contralateral cVNS elicits a larger LC response than contralateral taNS (Fig. 8, D, E). Interestingly, the largest IC responses are evoked by low intensity cVNS, indicating a U-shape relationship (Fig. 8, F). Regarding LC latency, increasing cVNS pulsing frequency (Fig. 9, B) and stimulus intensity (Fig. 9, F) is associated with faster LC responses, whereas no parameters alter LC latency in taNS (Fig. 9). The beta estimates for the EC amplitude and latency are reported in Tables S6, S7.

**Figure 8.**
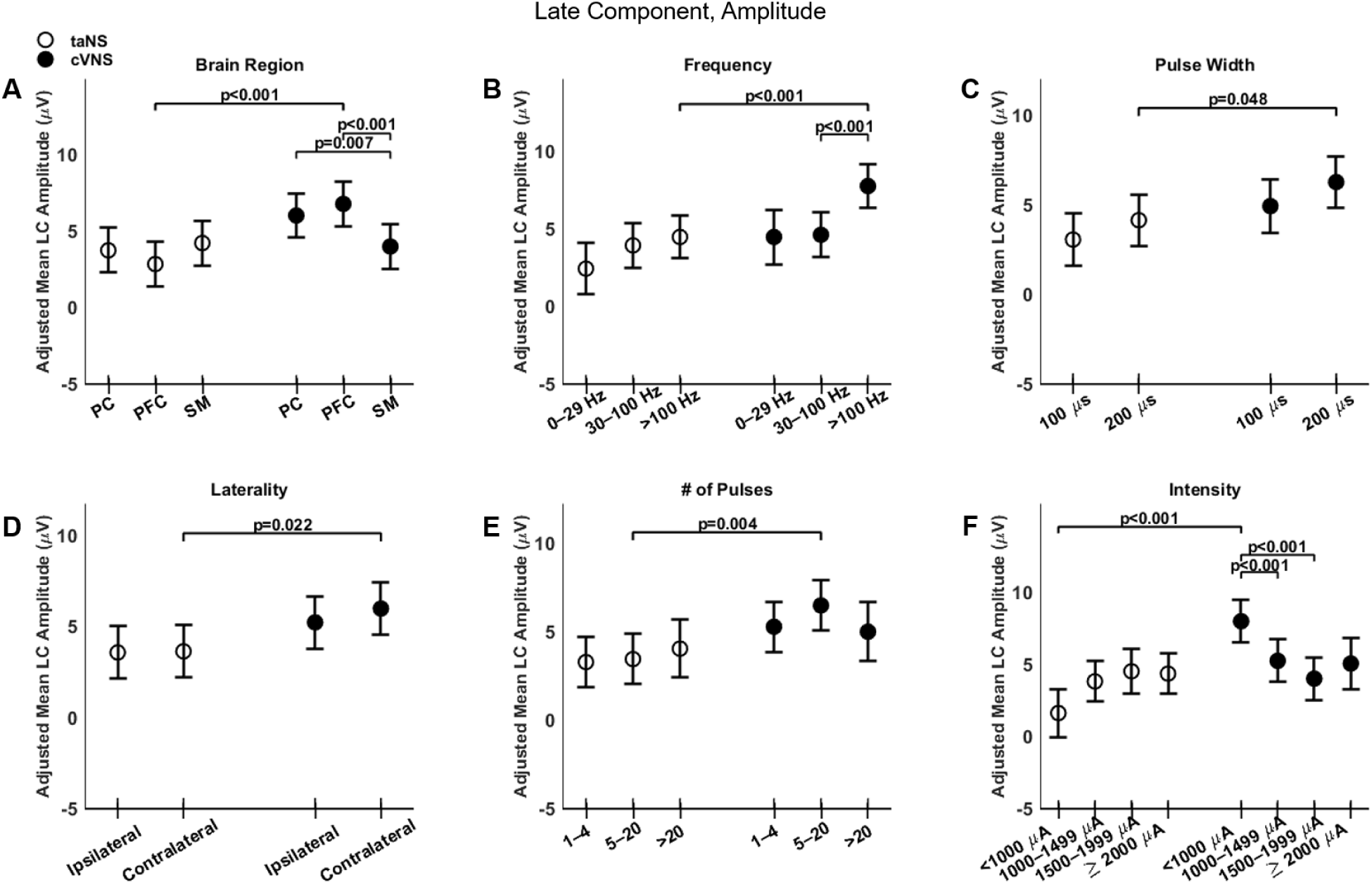
Effects of stimulation parameters on the amplitude of the late component. (A-F): Same as Fig. 4, but for the amplitude of the LC.

**Figure 9.**
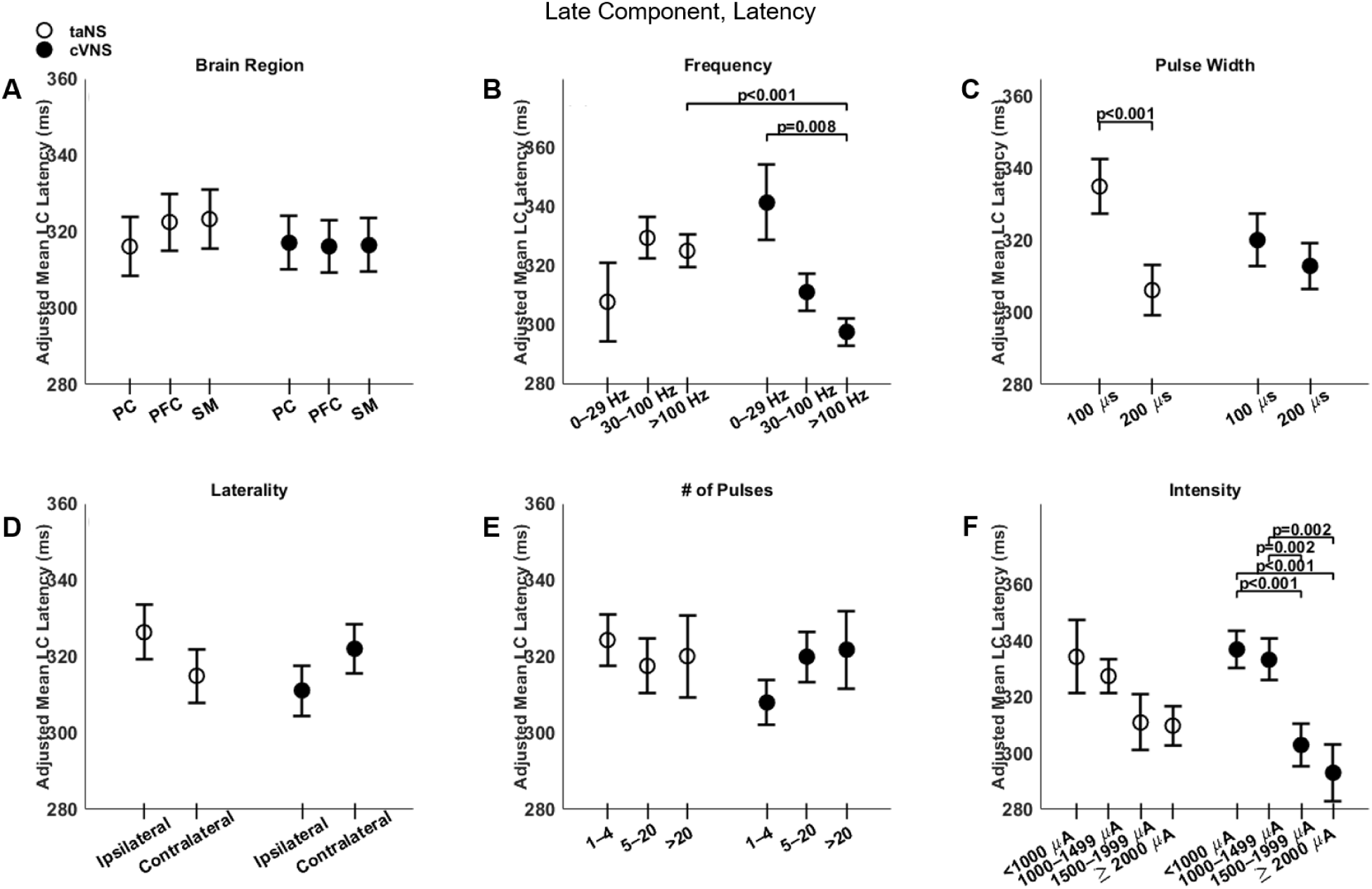
Effects of stimulation parameters on the latency of the late component (LC) of VEPs. (A-F): Same as Fig. 4, but for the latency of the LC.

## Discussion

The present report provides a descriptive, parametric characterization of cVNS- vs. taNS-elicited cortical VEPs recorded from six awake nonhuman primates. Our results reveal distinct similarities and differences between cVNS and taNS in terms of eliciting cortical responses, as well as differential effects of stimulation parameters on the amplitude and latency of VEP waveforms.

### Characterization and neural sources of cortical VEPs

We characterized cortical VEPs by measuring the amplitude and latency of (up to) 3 components: early, intermediate and late (Fig. 2, A). About 75% of VEPs have 1 or more components, whereas in 25% no VEP components are detected (Fig. 2, B). The choice of the latency windows for the 3 components was based on the distribution of latencies of detected waves from all recorded VEPs (Fig. 2, C) and is consistent with previous findings from characterization of VEPs in nonhuman primates engaged in different behavioral states [33]. We chose to use the absolute value of the amplitude of detected waves in our analysis and discard the polarity (negative or positive), as polarity of evoked potentials largely depends on the locations of the recording electrodes relative to subcortical and cortical generators of electrical potentials, and we did not have access to these relative locations in our subjects. In addition, we noticed that positive and negative amplitudes were almost equally represented within each of the 3 components (Suppl. Fig. S3), so there is small risk for amplitude polarity differences significantly confounding our analysis.

Cortical VEPs primarily reflect transient, time-locked effects of activation of ascending vagal pathways on ongoing cortical activity, rather than being a direct measure of neuronal responses in subcortical and cortical areas engaged by stimulation. With that said, and although cortical VEPs have been studied relatively sparingly [39]-[43], we can infer potential subcortical and cortical correlates of VEPs from neuroanatomical and neurophysiological studies on ascending vagal pathways, e.g., [43]-[47], as well as studies on the neural underpinnings of different components of sensory evoked potentials, e.g., [48]-[50]. The early component of VEPs (EC; 30-100 ms post-stimulus), likely reflects cortical potentials generated by relatively oligosynaptic, ascending pathways projecting to the cortex through the sensory thalamus [51], e.g., somatosensory fibers from pharynx and larynx, that would be activated by cVNS, or from the skin of the auricle, that would be activated by taNS [49], [52]. The intermediate VEP component (IC; 101-200 ms post-stimulus) potentially reflects activation of polysynaptic ascending pathways, starting at the brainstem nucleus tractus solitarius and the pontine parabrachial nucleus, through the mediodorsal or intralaminar thalamus [53], to insular and anterior cingulate cortex (ACC), as well as to prefrontal, frontal and fronto-parietal areas [47], [54].In addition, the IC could reflect cortical effects of activation of neuromodulatory projections to the cortex, either noradrenergic, through the locus coeruleus, or cholinergic, through basal forebrain nuclei [44], [55]-[56]. In principle, these pathways could be engaged by activation of either the sensory vagus nerve or of the auricular/trigeminal innervation of the auricle, even though at different levels. Finally, the late VEP component (LC; 201-500 ms post-stimulus) could reflects slower components of the same neuromodulatory inputs to the cortex, delayed and sustained cortical activation arising from limbic areas and the hypothalamus, and cortical activity arising from higher order thalamocortical relays and secondary cortico-cortical projections of the insula and ACC to prefrontal, premotor and other higher cortical areas [33], [53], [57]-[58].

### Differences and similarities between cVNS- vs. taNS-elicited VEPs

Overall, cVNS elicits larger cortical VEPs compared to taNS (Fig. 3, B; Table 4). This finding aligns with prior studies suggesting that cVNS provides more direct and robust activation of vagal afferents due to the proximity of the stimulus to the majority of vagus nerve fibers and the absence of intermediate tissue barriers [35],[59]. In contrast, taNS likely activate fewer sensory fibers, because of the relatively small number of auricular cutaneous neurons [60].

Larger cVNS-elicited VEPs, compared to taNS, are observed in PFC and PC areas, with regard to EC and LC (Fig. 4, A; 8, A), and in all cortical areas with regard to IC (Fig. 6, A), both ipsilateral and contralateral in both ipsilateral and contralateral to the side of stimulation (Fig. 4, D; 6, D; 8, D). The fact that cVNS generates larger early, intermediate and late VEP components across widespread cortical areas in both hemispheres suggests that cVNS more robustly activates oligosynaptic and polysynaptic ascending vagal pathways, bilaterally. Importantly, cVNS engages the PFC in a distinct manner compared to other cortical areas: it generates largest ECs and LCs, and smallest ICs. In contrast, taNS-elicited ICs are larger and faster in SM areas, but no area-specific responses in the other components (Fig. 4, A; 6, A; 8, A). The differences in distribution of cortical VEPs between the 2 stimulation modalities suggest that cVNS and taNS differentially engage cortical projections of ascending vagal pathways, possibly because they activate different first-order sensory neurons (visceral vs. cutaneous).

We found that cVNS-elicited ECs are slower than taNS-elicited ECs, by a relatively wide margin, of over 5 ms (Fig. 3, C). The effect is primarily driven by contralateral cVNS protocols (Fig. 5, D). One of the reasons behind this difference may be the longer distance afferent volleys travel when elicited from the cervical region (cVNS) vs. from the auricular region (taNS). Also, in addition to afferent fibers, cVNS activates efferent laryngeal fibers [61], triggering laryngeal muscle contractions, followed by secondary, afferent volleys in somatosensory fibers of the pharynx and larynx and in proprioceptive fibers of laryngeal muscles and joints [62], contributing to longer EC latencies. This is consistent with contralateral cVNS-elicited ECs being slower than taNS-elicited ECs (Fig. 5, D), as laryngeal and pharyngeal somatosensory pathways decussate to the opposite side before joining the medial lemniscus on their way to the thalamus. Latencies of ICs and LCs are similar between cVNS and taNS (Fig. 7, 9).

### Parameter-dependent effects on taNS-elicited VEPs

Different levels of stimulation parameters have distinct effects on taNS-elicited VEPs. taNS-elicited VEPs are no different between ipsilateral and contralateral sites, only exception being the latency of the EC, with contralateral taNS producing faster responses (Fig. 5, D). This suggests that the choice of left vs. right taNS may have limited impact on evoked brain responses. Higher taNS frequencies and intensities elicit larger ECs (Fig. 4, B, F) and ICs (Fig. 6, B, F), whereas LC amplitude is relatively insensitive to stimulation parameters (Fig. 6). Latencies of ICs and LCs tend to be faster at higher taNS intensities and longer pulse width, but the trend does not always reach statistical significance (Fig. 7, F; 9, F). These results are consistent with prior reports of positive correlations between taNS dose (frequency and/or intensity) with markers of brain or autonomic activity [41], [63]-[72], even though in some cases no such correlations were found [73]-[75]. Our findings suggest that such dose-response relationships may arise from differential engagement of ascending vagal afferent pathways, related to ECs and ICs, rather than from direct effects on higher order thalamocortical and cortico-cortical communication, related to LCs; this could be relevant to the growing use of taNS to improve cognitive and behavioral function in health and disease [13], [76]-[78]. Also, given the relatively small effects on amplitudes and latencies of VEPs, differential engagement of brain networks by taNS of varying parameters may be too subtle to be captured with non-specific measurements of autonomic function or even with neuroimaging methods with limited temporal resolution; this could be relevant to taNS optimization studies [34].

Using data from a relatively small number of experiments, we compared VEPs elicited by taNS and by trigeminal (ear lobe) stimulation, a site frequently used in studies as control condition (taCON). taCON elicits larger ECs, ICs and LCs, and faster ICs compared to taNS (Suppl. Fig. S4, A-C). However, the number of protocols tested for taCON was significantly smaller than for taNS (112 versus 2114). In addition, the ICC for the amplitude comparisons is large, indicating that individual differences among animals account for a significant portion of the observed variance (Suppl. Table S8). Despite these limitations, it is evident that stimulation of the ear lobe engages robustly both fast and slower ascending sensory pathways, resulting in cortical activation patterns that may be partially overlapping with those in response to taNS.

### Parameter-dependent effects on cVNS-elicited VEPs

cVNS-elicited VEPs exhibit specific parameter-dependent effects. Overall, ipsilateral cVNS elicits larger ECs than contralateral cVNS (Fig. 4, D). Even though the effect is of moderate size, it may be relevant to the use of chronic cVNS in rehabilitation after unilateral stroke [79]. The amplitude of EC and IC responses is strongly positively correlated with increasing pulsing frequency and stimulation intensity (Fig. 4, 6, 8), in agreement with prior animal work documenting these effects on subcortical and cortical responses, neuronal activity and neurotransmitter release, both in animals [43], [80]-[83] and in humans [29], [84]-[88]. In our study, we did not observe many U- or inverse U-shaped relationships, previously documented in both acute markers of vagal pathway engagement and in more sustained, cortical plasticity outcomes, e.g., [83],[89]-[90]. Notable exceptions are the effect of train duration on the amplitude of the EC (Fig. 4, E), consistent with the observation that longer VNS trains lead to adaption of neuronal responses [43], and the effect of intensity on the amplitude of the LC (Fig. 8, F). Given that VEPs are indirect markers of cortical network engagement, our results suggest that some of the non-monotonic relationships between stimulus parameters and direct markers of brain activation may arise from non-linear interactions between stimulus-triggered activation of ascending vagal pathways and other local and/or distributed neuronal networks that further shape neuronal responses to neurostimulation, e.g., [33].

### Study limitations

While our study provides valuable insights into the differential effects of cVNS and taNS, it has several limitations. First, the use of an animal model limits direct translation to human physiology, although the similarities in vagal-cortical pathways between humans and nonhuman primates support the relevance of our findings [96]. Second, the use of a cortical implant limited our findings to cortical VEPs, allowing us to indirectly and partially infer the effects of cVNS and taNS on subcortical sites along the ascending vagal pathways [97]. Brain mechanisms shaping parameter-specific effects, particularly the role of neural plasticity [98] and longitudinal changes in the electrode-tissue interface [99], warrant further exploration. Finally, even though parametric studies, such as this one, provide valuable information for optimization of neuromodulation therapies, our study does not provide direct evidence for which stimulus parameters may be more effective or what features of evoked brain responses may be predictive of a clinical response to VNS.

## Conclusion

In summary, our study demonstrates that cVNS and taNS elicit distinct patterns of cortical activation, with cVNS producing larger and faster VEPs compared to taNS, in most sampled cortical areas. We identify key stimulation parameters that influence VEP characteristics, providing a foundation for optimizing neuromodulation protocols. These findings contribute to a growing body of literature on how VNS affects brain function and highlight the potential for tailored approaches to enhance therapeutic efficacy.

## Acknowledgments

We thank Larry Shupe for his technical support. Part of this work was supported by grants DARPA HR0011-17-2-0025 to SZ and EF, and by NIH 1R01NS136685-01A1 to SZ.

## Data Availability

The raw Vegal Evoked Potentials VEPs with the processed data used for the LLM generated for this study are available in the Figshare repository: ‘VEPs non-human primates: Rembado I., Ravan M, et al. (2026), Cortical potentials evoked by stimulation of cervical vagus vs. auricular nerve: A comparative, parametric study in nonhuman primates (https://doi.org/10.6084/m9.figshare.30728222).

