## Supplementary Material for "Cortical potentials evoked by stimulation of cervical vagus vs. auricular nerve: a comparative, parametric study in nonhuman primates"

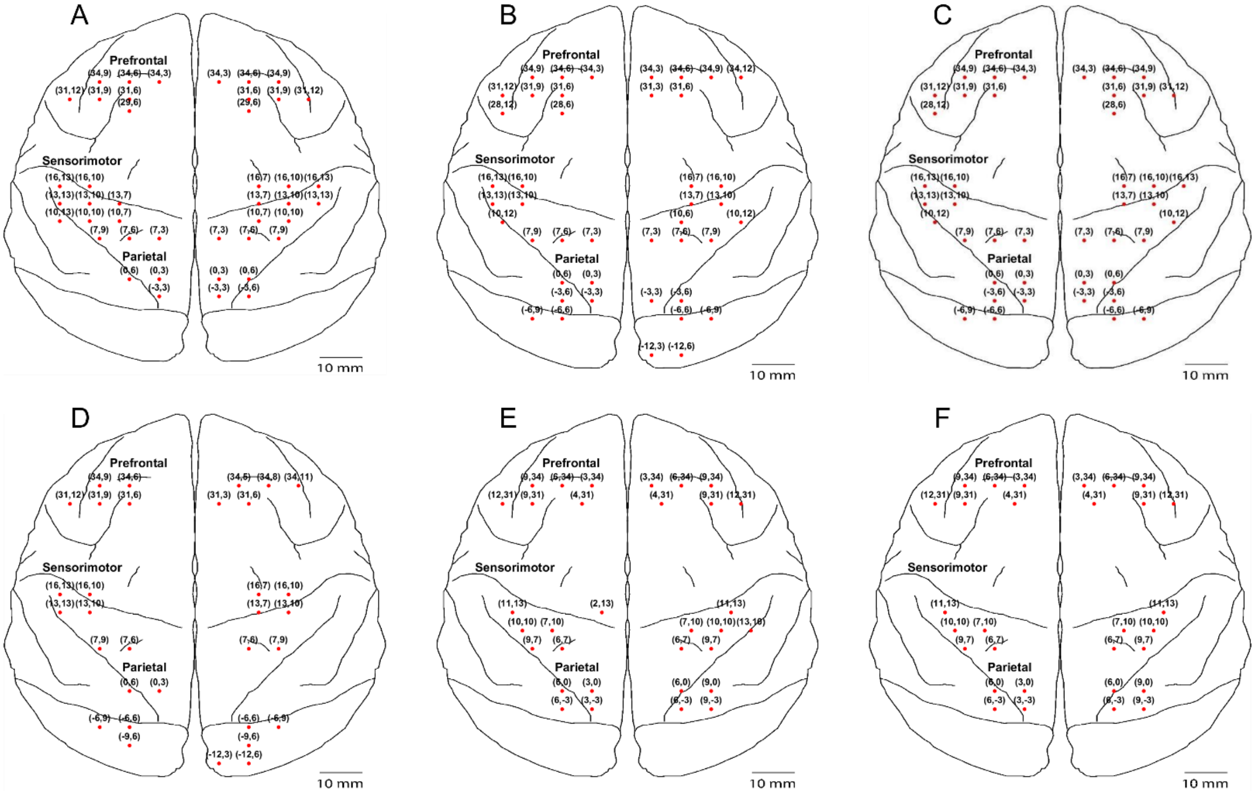


**Supplementary Figure S1.** Cortical locations of implanted electrodes on the cortical map for Bean (A), Hugo (B), Gomez (C), Inigo (D), Draco (E), and Zhivago (F). For each site, the numbers in parentheses indicate the stereotaxic coordinates: medial-lateral (distance from midline in mm), and anterior-posterior (distance from anterior commissure, in mm).

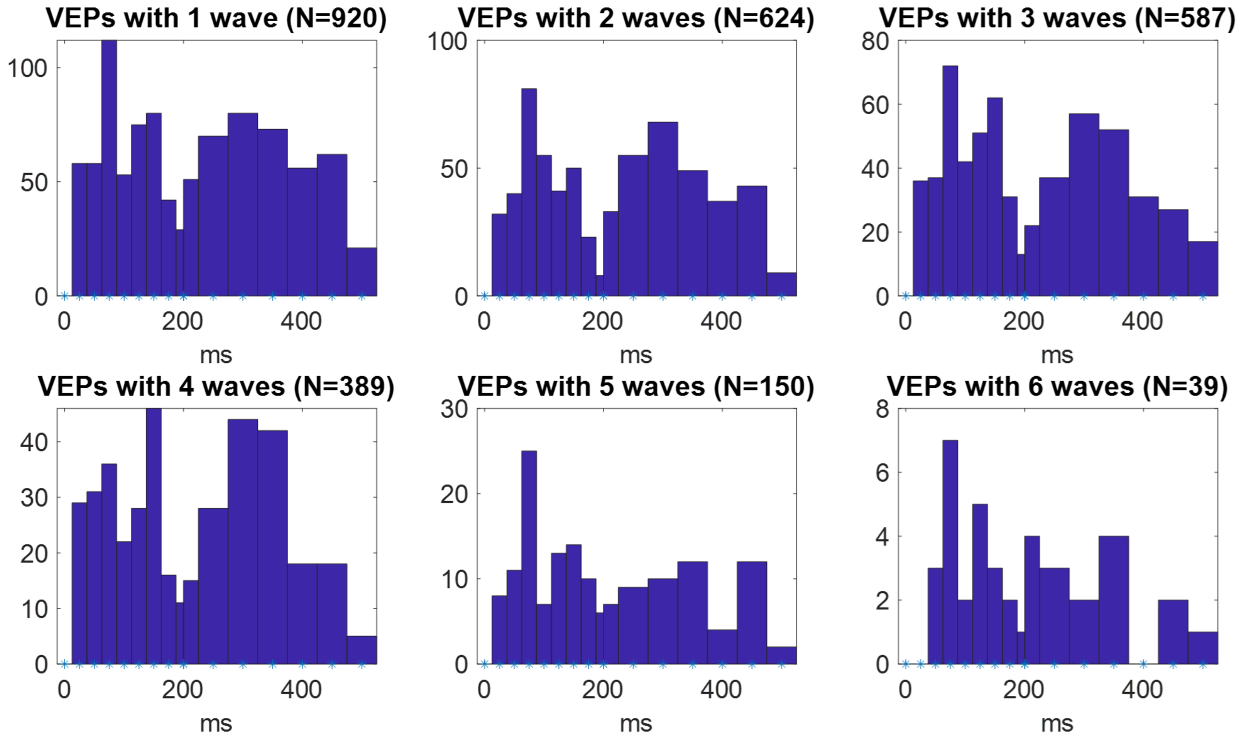


**Supplementary Figure S2.** Latency distributions of detected waves in subsets of VEPs with 1, 2, 3, etc waves.

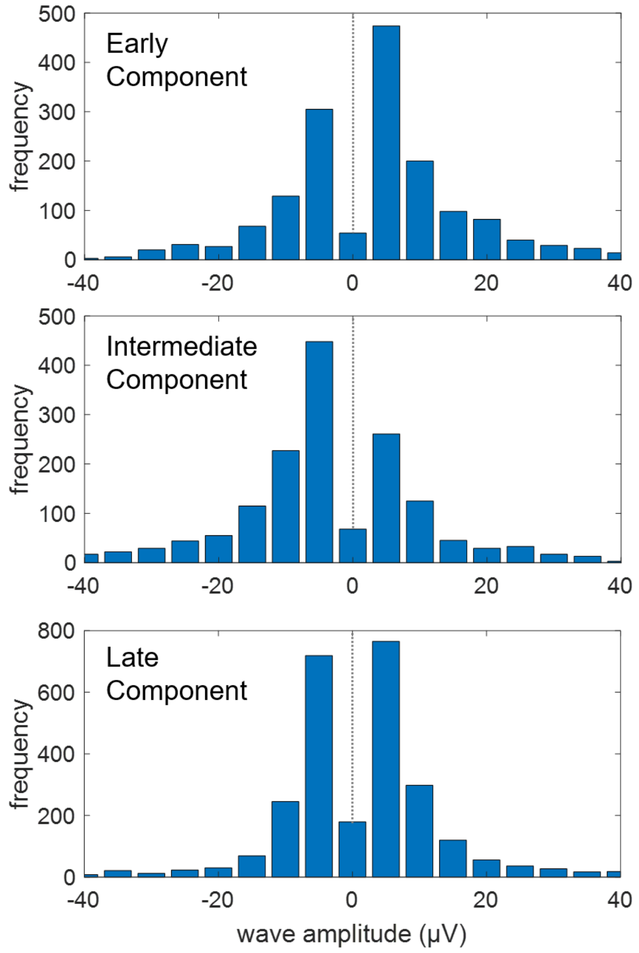


**Supplementary Figure S3.** Amplitude distributions of detected waves classified as early (top), intermediate (middle) or late components (bottom).

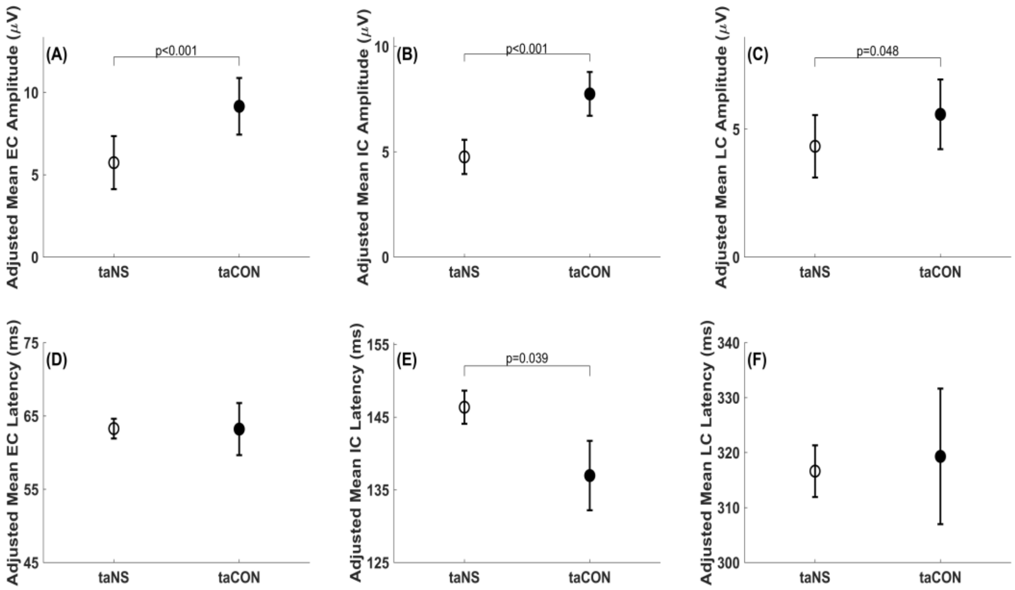


**Supplementary Figure S4.** Overall comparison of VEPs between taNS and trans-auricular control stimulation (taCON), applied to the earlobe. (A-C) Adjusted means for EC, IC and LC amplitudes. (D-F) Adjusted means for EC, IC and LC latencies.

**Supplementary Table S1.** Breakdown of the stimulation protocols used in each monkey

| Monkeys | | | | | N. Pulses group | Intens-ity group | Freq-ency group | Pulse width | Stim site | N. trials |
|---|---|---|---|---|---|---|---|---|---|---|
| Bean | Gomez | Hugo | Inigo | | 5–20 | 1000–1499 | 30–100 | 100 | taNS | 67 |
| Zhivago | | | | | 5–20 | ≥2000 | >100 | 200 | cvns | 45 |
| Bean | Gomez | Hugo | Inigo | | 5–20 | 1000–1499 | 30–100 | 200 | taNS | 44 |
| Bean | Draco | Hugo | Inigo | Zhivago | 5–20 | 1000–1499 | >100 | 200 | taNS | 41 |
| Bean | Gomez | Hugo | | | 5–20 | ≥2000 | 30–100 | 100 | taNS | 40 |
| Draco | Zhivago | | | | 5–20 | 1000–1499 | >100 | 200 | cvns | 39 |
| Draco | Inigo | | | | 5–20 | <1000 | 30–100 | 200 | cvns | 36 |
| Draco | Inigo | | | | 5–20 | <1000 | >100 | 200 | cvns | 34 |
| Bean | Draco | Hugo | Zhivago | | 5–20 | ≥2000 | >100 | 200 | taNS | 32 |
| Bean | Gomez | Hugo | Inigo | | >20 | 1000–1499 | >100 | 100 | taNS | 31 |
| Bean | Draco | Gomez | Hugo | | 5–20 | <1000 | >100 | 200 | taNS | 27 |
| Draco | Gomez | Inigo | | | 5–20 | 1000–1499 | 30–100 | 200 | cvns | 25 |
| Draco | Zhivago | | | | 5–20 | 1500–1999 | >100 | 200 | cvns | 24 |

| | | | | | | | | | | |
|---|---|---|---|---|---|---|---|---|---|---|
| Inigo | | | | | 5–20 | <1000 | 30–100 | 100 | cvns | 23 |
| Bean | Hugo | Zhivago | | | 5–20 | 1500–1999 | >100 | 200 | taNS | 23 |
| Gomez | | | | | 5–20 | 1000–1499 | 30–100 | 100 | cvns | 21 |
| Bean | Gomez | Hugo | | | >20 | ≥2000 | >100 | 100 | taNS | 16 |
| Draco | Gomez | Zhivago | | | 5–20 | 1500–1999 | 30–100 | 200 | cvns | 15 |
| Zhivago | | | | | 5–20 | ≥2000 | 30–100 | 200 | cvns | 14 |
| Gomez | Inigo | | | | >20 | 1000–1499 | >100 | 200 | taNS | 14 |
| Gomez | | | | | 5–20 | 1500–1999 | 30–100 | 100 | cvns | 13 |
| Bean | Gomez | Hugo | | | 1–4 | ≥2000 | 0–29 | 200 | taNS | 11 |
| Inigo | | | | | >20 | <1000 | >100 | 200 | cvns | 11 |
| Gomez | Inigo | | | | 5–20 | 1000–1499 | >100 | 100 | cvns | 10 |
| Bean | Gomez | Hugo | | | 5–20 | ≥2000 | 30–100 | 200 | taNS | 9 |
| Bean | Draco | Hugo | Zhivago | | 5–20 | ≥2000 | >100 | 200 | ta Control | 9 |
| Inigo | | | | | >20 | <1000 | >100 | 100 | cvns | 9 |
| Bean | Zhivago | | | | 5–20 | 1500–1999 | >100 | 200 | ta Control | 8 |
| Hugo | Inigo | | | | 1–4 | 1000–1499 | 0–29 | 100 | taNS | 7 |
| Bean | Gomez | Hugo | Inigo | | 1–4 | 1000–1499 | 0–29 | 200 | taNS | 7 |
| Bean | Gomez | Hugo | | | 5–20 | <1000 | 30–100 | 200 | taNS | 7 |
| Bean | Inigo | | | | 5–20 | 1000–1499 | >100 | 100 | taNS | 7 |
| Bean | | | | | 5–20 | ≥2000 | >100 | 100 | taNS | 7 |
| Gomez | Inigo | | | | 1–4 | <1000 | 0–29 | 100 | cvns | 6 |
| Gomez | Inigo | | | | 1–4 | 1500–1999 | 0–29 | 200 | cvns | 6 |
| Bean | Hugo | Inigo | | | 5–20 | 1500–1999 | 30–100 | 200 | taNS | 6 |
| Bean | | | | | 5–20 | 1500–1999 | >100 | 100 | taNS | 6 |
| Gomez | | | | | >20 | 1000–1499 | >100 | 100 | cvns | 6 |
| Gomez | | | | | >20 | 1500–1999 | >100 | 100 | cvns | 6 |
| Bean | Draco | | | | 5–20 | 1000–1499 | >100 | 200 | ta Control | 5 |
| Bean | Hugo | Inigo | | | 1–4 | 1500–1999 | 0–29 | 200 | taNS | 4 |
| Hugo | | | | | 1–4 | ≥2000 | 0–29 | 100 | taNS | 4 |
| Inigo | | | | | 1–4 | <1000 | 0–29 | 200 | cvns | 3 |

| | | | | | | | | | | |
|---|---|---|---|---|---|---|---|---|---|---|
| Gomez | Inigo | | | | 5–20 | <1000 | >100 | 100 | cvns | 3 |
| Gomez | | | | | 5–20 | 1500–1999 | >100 | 100 | cvns | 3 |
| Gomez | | | | | >20 | 1000–1499 | >100 | 200 | cvns | 3 |
| Gomez | | | | | 1–4 | 1000–1499 | 0–29 | 100 | cvns | 2 |
| Gomez | | | | | 1–4 | 1500–1999 | 0–29 | 100 | cvns | 2 |
| Bean | | | | | 5–20 | 1500–1999 | 30–100 | 100 | taNS | 2 |
| Bean | | | | | 5–20 | 1500–1999 | 30–100 | 200 | ta Control | 2 |
| Gomez | | | | | >20 | ≥2000 | >100 | 200 | taNS | 2 |
| Bean | | | | | 5–20 | 1000–1499 | 30–100 | 200 | ta Control | 1 |
| Bean | | | | | 5–20 | ≥2000 | 30–100 | 200 | ta Control | 1 |
| Gomez | | | | | 5–20 | ≥2000 | >100 | 100 | cvns | 1 |

**Supplementary Table S2.** LMM beta estimates for EC amplitude

| Effect | | | Beta Estimate | Standard Error | Lower CL | Upper CL | Pr > \|t\| |
|---|---|---|---|---|---|---|---|
| **Intercept** | Variable | Interaction variable | 2.20 | 2.17 | -2.05 | 6.46 | 0.3568 |
| **Stimulation site** | cVNS | | -1.36 | 2.08 | -5.44 | 2.72 | 0.5134 |
| | taNS | | 0.00 | . | | | . |
| **Brain region** | PC | | -0.79 | 0.67 | -2.09 | 0.51 | 0.2355 |
| | PFC | | -1.29 | 0.65 | -2.56 | -0.02 | 0.0458 |
| | SM | | 0.00 | . | | | . |
| **# of pulses** | Intermediate [5-20) | | -0.12 | 0.64 | -1.36 | 1.13 | 0.8544 |
| | Long 20+ | | -1.77 | 1.11 | -3.94 | 0.40 | 0.1093 |
| | Short [1-4] | | 0.00 | . | | | . |
| **Intensity** | 2000+ | | 3.71 | 1.14 | 1.47 | 5.95 | 0.0012 |
| | [1000-1500) | | 1.88 | 1.10 | -0.29 | 4.04 | 0.0893 |
| | [1500-2000) | | 2.11 | 1.32 | -0.48 | 4.70 | 0.1099 |
| | <1000 | | 0.00 | . | | | . |
| **Pulse width** | 100 | | -1.53 | 0.62 | -2.76 | -0.31 | 0.0142 |
| | 200 | | 0.00 | . | | | . |
| **Frequency** | High 100+ Hz | | 3.38 | 1.20 | 1.03 | 5.73 | 0.0048 |
| | Medium [30-100 Hz] | | 0.93 | 1.05 | -1.13 | 2.99 | 0.3756 |
| | Low [0-30 Hz) | | 0.00 | . | | | . |

| | | | | | | | |
|---|---|---|---|---|---|---|---|
| **Laterality** | Contralateral | | 0.08 | 0.53 | -0.97 | 1.13 | 0.885 |
| | Ipsilateral | | 0.00 | . | | | . |
| **Stimulation site * Brain region** | cVNS | PC | 5.34 | 0.95 | 3.47 | 7.21 | <.0001 |
| | cVNS | PFC | 6.02 | 0.94 | 4.18 | 7.86 | <.0001 |
| | cVNS | SM | 0.00 | . | | | . |
| | taNS | PC | 0.00 | . | | | . |
| | taNS | PFC | 0.00 | . | | | . |
| | taNS | SM | 0.00 | . | | | . |
| **Stimulation site * # of pulses** | cVNS | Intermediate [5-20) | 2.16 | 0.91 | 0.37 | 3.95 | 0.0183 |
| | cVNS | Long 20+ | -1.06 | 1.56 | -4.12 | 2.00 | 0.4964 |
| | cVNS | Short [1-4] | 0.00 | . | | | . |
| | taNS | Intermediate [5-20) | 0.00 | . | | | . |
| | taNS | Long 20+ | 0.00 | . | | | . |
| | taNS | Short [1-4] | 0.00 | . | | | . |
| **Stimulation site * Intensity** | cVNS | 2000+ | 1.26 | 1.82 | -2.30 | 4.83 | 0.4869 |
| | cVNS | [1000-1500) | 1.95 | 1.38 | -0.75 | 4.65 | 0.1576 |
| | cVNS | [1500-2000) | 0.62 | 1.58 | -2.49 | 3.73 | 0.6956 |
| | cVNS | <1000 | 0.00 | . | | | . |
| | taNS | 2000+ | 0.00 | . | | | . |
| | taNS | [1000-1500) | 0.00 | . | | | . |
| | taNS | [1500-2000) | 0.00 | . | | | . |
| | taNS | <1000 | 0.00 | . | | | . |
| **Stimulation site * Pulse width** | cVNS | 100 | -0.33 | 1.01 | -2.32 | 1.66 | 0.7437 |
| | cVNS | 200 | 0.00 | . | | | . |
| | taNS | 100 | 0.00 | . | | | . |
| | taNS | 200 | 0.00 | . | | | . |
| **Stimulation site * Frequency** | cVNS | High 100+ Hz | 2.14 | 1.81 | -1.41 | 5.68 | 0.2385 |
| | cVNS | Medium [30-100 Hz] | -1.43 | 1.68 | -4.72 | 1.86 | 0.3941 |
| | cVNS | Low [0-30 Hz) | 0.00 | . | | | . |
| | taNS | High 100+ Hz | 0.00 | . | | | . |
| | taNS | Medium [30-100 Hz] | 0.00 | . | | | . |
| | taNS | Low [0-30 Hz) | 0.00 | . | | | . |
| **Stimulation site * Laterality** | cVNS | Contralateral | -2.55 | 0.77 | -4.06 | -1.03 | 0.001 |
| | cVNS | Ipsilateral | 0.00 | . | | | . |

| | | | | | | | |
|---|---|---|---|---|---|---|---|
| | taNS | Contralateral | 0.00 | . | | | . |
| | taNS | Ipsilateral | 0.00 | . | | | . |

**Supplementary Table S3.** LMM beta estimate for the EC latency

| Effect | | | Beta Estimate | Standard Error | Lower CL | Upper CL | Pr > \|t\| |
|---|---|---|---|---|---|---|---|
| **Intercept** | | | 51.64 | 4.58 | 42.66 | 60.63 | <.0001 |
| **Stimulation site** | cVNS | | 11.37 | 5.80 | 0.00 | 22.75 | 0.0503 |
| | taNS | | 0.00 | . | | | . |
| **Brain region** | PC | | 3.23 | 1.65 | 0.00 | 6.46 | 0.0504 |
| | PFC | | 0.08 | 1.60 | -3.06 | 3.22 | 0.9602 |
| | SM | | 0.00 | . | | | . |
| **# of pulses** | Intermediate [5-20) | | 0.99 | 1.59 | -2.11 | 4.10 | 0.5306 |
| | Long 20+ | | -5.49 | 2.57 | -10.52 | -0.47 | 0.0324 |
| | Short [1-4] | | 0.00 | . | | | . |
| **Intensity** | 2000+ | | 2.47 | 3.32 | -4.04 | 8.98 | 0.4567 |
| | [1000-1500) | | 3.49 | 3.34 | -3.05 | 10.03 | 0.2956 |
| | [1500-2000) | | 4.82 | 3.61 | -2.26 | 11.89 | 0.1823 |
| | <1000 | | 0.00 | . | | | . |
| **Pulse width** | 100 | | 2.01 | 1.66 | -1.23 | 5.26 | 0.2243 |
| | 200 | | 0.00 | . | | | . |
| **Frequency** | High 100+ Hz | | 9.20 | 3.31 | 2.71 | 15.70 | 0.0055 |
| | Medium [30-100 Hz] | | 7.52 | 3.20 | 1.26 | 13.79 | 0.0187 |
| | Low [0-30 Hz) | | 0.00 | . | | | . |
| **Laterality** | Contralateral | | -4.00 | 1.34 | -6.62 | -1.38 | 0.0028 |
| | Ipsilateral | | 0.00 | . | | | . |
| **Stimulation site x Brain region** | cVNS | PC | 0.36 | 2.30 | -4.15 | 4.88 | 0.8744 |
| | cVNS | PFC | 2.05 | 2.22 | -2.29 | 6.40 | 0.3549 |
| | cVNS | SM | 0.00 | . | | | . |
| | taNS | PC | 0.00 | . | | | . |
| | taNS | PFC | 0.00 | . | | | . |
| | taNS | SM | 0.00 | . | | | . |
| **Stimulation site x # of pulses** | cVNS | Intermediate [5-20) | -2.16 | 2.21 | -6.48 | 2.16 | 0.3277 |
| | cVNS | Long 20+ | 1.33 | 3.50 | -5.53 | 8.19 | 0.7036 |
| | cVNS | Short [1-4] | 0.00 | . | | | . |
| | taNS | Intermediate [5-20) | 0.00 | . | | | . |
| | taNS | Long 20+ | 0.00 | . | | | . |
| | taNS | Short [1-4] | 0.00 | . | | | . |
| | cVNS | 2000+ | -2.82 | 4.06 | -10.78 | 5.14 | 0.4878 |

| | | | | | | | |
|---|---|---|---|---|---|---|---|
| Stimulation site x Intensity | cVNS | [1000-1500) | -5.63 | 3.91 | -13.28 | 2.03 | 0.1500 |
| | cVNS | [1500-2000) | -5.94 | 4.17 | -14.12 | 2.24 | 0.1548 |
| | cVNS | <1000 | 0.00 | . | | | . |
| | taNS | 2000+ | 0.00 | . | | | . |
| | taNS | [1000-1500) | 0.00 | . | | | . |
| | taNS | [1500-2000) | 0.00 | . | | | . |
| | taNS | <1000 | 0.00 | . | | | . |
| **Stimulation site x Pulse width** | cVNS | 100 | -5.82 | 2.45 | -10.63 | -1.02 | 0.0176 |
| | cVNS | 200 | 0.00 | . | | | . |
| | taNS | 100 | 0.00 | . | | | . |
| | taNS | 200 | 0.00 | . | | | . |
| **Stimulation site x Frequency** | cVNS | High 100+ Hz | 0.05 | 4.83 | -9.42 | 9.52 | 0.9913 |
| | cVNS | Medium [30-100 Hz] | -3.24 | 4.69 | -12.44 | 5.95 | 0.4892 |
| | cVNS | Low [0-30 Hz) | 0.00 | . | | | . |
| | taNS | High 100+ Hz | 0.00 | . | | | . |
| | taNS | Medium [30-100 Hz] | 0.00 | . | | | . |
| | taNS | Low [0-30 Hz) | 0.00 | . | | | . |
| **Stimulation site x Laterality** | cVNS | Contralateral | 3.71 | 1.84 | 0.11 | 7.31 | 0.0436 |
| | cVNS | Ipsilateral | 0.00 | . | | | . |
| | taNS | Contralateral | 0.00 | . | | | . |
| | taNS | Ipsilateral | 0.00 | . | | | . |

**Supplementary Table S4.** LMM beta estimates for IC amplitude

| Effect | | | Beta Estimate | Standard Error | Lower CL | Upper CL | Pr > \|t\| |
|---|---|---|---|---|---|---|---|
| **Intercept** | | | 0.21 | 1.59 | -2.91 | 3.32 | 0.9023 |
| **Stimulation site** | cVNS | | 2.52 | 1.95 | -1.31 | 6.35 | 0.1972 |
| | taNS | | 0.00 | . | | | . |
| **Brain region** | PC | | -1.35 | 0.63 | -2.58 | -0.12 | 0.0316 |
| | PFC | | -1.91 | 0.61 | -3.11 | -0.72 | 0.0017 |
| | SM | | 0.00 | . | | | . |
| **# of pulses** | Intermediate [5-20) | | 0.41 | 0.60 | -0.76 | 1.59 | 0.4876 |
| | Long 20+ | | -1.22 | 1.04 | -3.26 | 0.82 | 0.2412 |
| | Short [1-4] | | 0.00 | . | | | . |
| **Intensity** | 2000+ | | 4.36 | 1.07 | 2.25 | 6.47 | <.0001 |
| | [1000-1500) | | 2.24 | 1.04 | 0.20 | 4.28 | 0.0316 |
| | [1500-2000) | | 3.17 | 1.24 | 0.73 | 5.61 | 0.0108 |
| | <1000 | | 0.00 | . | | | . |

| | | | | | | | |
|---|---|---|---|---|---|---|---|
| **Pulse width** | 100 | | -1.51 | 0.59 | -2.66 | -0.36 | 0.0104 |
| | 200 | | 0.00 | . | | | . |
| **Frequency** | High 100+ Hz | | 4.03 | 1.13 | 1.82 | 6.23 | 0.0003 |
| | Medium [30-100 Hz] | | 0.93 | 0.99 | -1.01 | 2.87 | 0.3453 |
| | Low [0-30 Hz) | | 0.00 | . | | | . |
| **Laterality** | Contralateral | | 0.25 | 0.50 | -0.74 | 1.24 | 0.6232 |
| | Ipsilateral | | 0.00 | . | | | . |
| **Stimulation site * Brain region** | cVNS | PC | 2.20 | 0.90 | 0.44 | 3.97 | 0.0145 |
| | cVNS | PFC | -0.20 | 0.89 | -1.94 | 1.54 | 0.8196 |
| | cVNS | SM | 0.00 | . | | | . |
| | taNS | PC | 0.00 | . | | | . |
| | taNS | PFC | 0.00 | . | | | . |
| | taNS | SM | 0.00 | . | | | . |
| **Stimulation site * # of pulses** | cVNS | Intermediate [5-20) | -0.36 | 0.86 | -2.05 | 1.33 | 0.6786 |
| | cVNS | Long 20+ | -1.11 | 1.47 | -3.99 | 1.77 | 0.4488 |
| | cVNS | Short [1-4] | 0.00 | . | | | . |
| | taNS | Intermediate [5-20) | 0.00 | . | | | . |
| | taNS | Long 20+ | 0.00 | . | | | . |
| | taNS | Short [1-4] | 0.00 | . | | | . |
| **Stimulation site * Intensity** | cVNS | 2000+ | 3.62 | 1.67 | 0.35 | 6.89 | 0.0302 |
| | cVNS | [1000-1500) | 0.00 | 1.30 | -2.54 | 2.54 | 0.9985 |
| | cVNS | [1500-2000) | 1.31 | 1.49 | -1.61 | 4.24 | 0.3786 |
| | cVNS | <1000 | 0.00 | . | | | . |
| | taNS | 2000+ | 0.00 | . | | | . |
| | taNS | [1000-1500) | 0.00 | . | | | . |
| | taNS | [1500-2000) | 0.00 | . | | | . |
| | taNS | <1000 | 0.00 | . | | | . |
| **Stimulation site * Pulse width** | cVNS | 100 | -0.61 | 0.95 | -2.47 | 1.25 | 0.5184 |
| | cVNS | 200 | 0.00 | . | | | . |
| | taNS | 100 | 0.00 | . | | | . |
| | taNS | 200 | 0.00 | . | | | . |
| **Stimulation site * Frequency** | cVNS | High 100+ Hz | 2.95 | 1.71 | -0.39 | 6.30 | 0.0836 |
| | cVNS | Medium [30-100 Hz] | -0.50 | 1.58 | -3.60 | 2.61 | 0.7542 |
| | cVNS | Low [0-30 Hz) | 0.00 | . | | | . |
| | taNS | High 100+ Hz | 0.00 | . | | | . |
| | taNS | Medium [30-100 Hz] | 0.00 | . | | | . |
| | taNS | Low [0-30 Hz) | 0.00 | . | | | . |
| | cVNS | Contralateral | -0.94 | 0.73 | -2.36 | 0.49 | 0.1967 |

| | | | | | | | |
|---|---|---|---|---|---|---|---|
| Stimulation site * Laterality | cVNS | Ipsilateral | 0.00 | . | | | . |
| | taNS | Contralateral | 0.00 | . | | | . |
| | taNS | Ipsilateral | 0.00 | . | | | . |

**Supplementary Table S5.** LMM beta estimates for IC latency

| Effect | | | Beta Estimate | Standard Error | Lower CL | Upper CL | Pr > \|t\| |
|---|---|---|---|---|---|---|---|
| **Intercept** | | | 139.99 | 6.44 | 127.38 | 152.60 | <.0001 |
| **Stimulation site** | cVNS | | -4.98 | 7.55 | -19.78 | 9.83 | 0.5102 |
| | taNS | | 0.00 | . | | | . |
| **Brain region** | PC | | 3.97 | 2.31 | -0.56 | 8.49 | 0.0859 |
| | PFC | | 9.37 | 2.17 | 5.12 | 13.62 | <.0001 |
| | SM | | 0.00 | . | | | . |
| **# of pulses** | Intermediate [5-20) | | 3.26 | 2.20 | -1.05 | 7.57 | 0.1386 |
| | Long 20+ | | 3.95 | 3.46 | -2.82 | 10.73 | 0.2529 |
| | Short [1-4] | | 0.00 | . | | | . |
| **Intensity** | 2000+ | | -3.10 | 4.43 | -11.79 | 5.58 | 0.4838 |
| | [1000-1500) | | 1.92 | 4.47 | -6.84 | 10.67 | 0.6682 |
| | [1500-2000) | | -7.64 | 4.82 | -17.08 | 1.80 | 0.1131 |
| | <1000 | | 0.00 | . | | | . |
| **Pulse width** | 100 | | 0.86 | 2.24 | -3.54 | 5.25 | 0.7016 |
| | 200 | | 0.00 | . | | | . |
| **Frequency** | High 100+ Hz | | 0.43 | 4.42 | -8.24 | 9.10 | 0.9224 |
| | Medium [30-100 Hz] | | 5.78 | 4.26 | -2.57 | 14.12 | 0.1752 |
| | Low [0-30 Hz) | | 0.00 | . | | | . |
| **Laterality** | Contralateral | | 1.78 | 1.85 | -1.84 | 5.40 | 0.3363 |
| | Ipsilateral | | 0.00 | . | | | . |
| **Stimulation site * Brain region** | cVNS | PC | 1.40 | 3.15 | -4.77 | 7.57 | 0.6558 |
| | cVNS | PFC | -4.50 | 3.05 | -10.49 | 1.49 | 0.1408 |
| | cVNS | SM | 0.00 | . | | | . |
| | taNS | PC | 0.00 | . | | | . |
| | taNS | PFC | 0.00 | . | | | . |
| | taNS | SM | 0.00 | . | | | . |
| **Stimulation site * # of pulses** | cVNS | Intermediate [5-20) | 0.20 | 3.05 | -5.77 | 6.18 | 0.9467 |
| | cVNS | Long 20+ | 10.08 | 4.65 | 0.97 | 19.20 | 0.0303 |
| | cVNS | Short [1-4] | 0.00 | . | | | . |
| | taNS | Intermediate [5-20) | 0.00 | . | | | . |
| | taNS | Long 20+ | 0.00 | . | | | . |

| | | | | | | | |
|---|---|---|---|---|---|---|---|
| | taNS | Short [1-4] | 0.00 | . | | | . |
| **Stimulation site * Intensity** | cVNS | 2000+ | 3.15 | 5.88 | -8.36 | 14.67 | 0.5914 |
| | cVNS | [1000-1500) | -3.24 | 5.40 | -13.82 | 7.34 | 0.5481 |
| | cVNS | [1500-2000) | 7.61 | 5.72 | -3.61 | 18.83 | 0.1838 |
| | cVNS | <1000 | 0.00 | . | | | . |
| | taNS | 2000+ | 0.00 | . | | | . |
| | taNS | [1000-1500) | 0.00 | . | | | . |
| | taNS | [1500-2000) | 0.00 | . | | | . |
| | taNS | <1000 | 0.00 | . | | | . |
| **Stimulation site * Pulse width** | cVNS | 100 | 2.73 | 3.31 | -3.75 | 9.22 | 0.409 |
| | cVNS | 200 | 0.00 | . | | | . |
| | taNS | 100 | 0.00 | . | | | . |
| | taNS | 200 | 0.00 | . | | | . |
| **Stimulation site * Frequency** | cVNS | High 100+ Hz | -4.48 | 6.13 | -16.51 | 7.54 | 0.4648 |
| | cVNS | Medium [30-100 Hz] | 1.18 | 5.89 | -10.36 | 12.72 | 0.8409 |
| | cVNS | Low [0-30 Hz) | 0.00 | . | | | . |
| | taNS | High 100+ Hz | 0.00 | . | | | . |
| | taNS | Medium [30-100 Hz] | 0.00 | . | | | . |
| | taNS | Low [0-30 Hz) | 0.00 | . | | | . |
| **Stimulation site * Laterality** | cVNS | Contralateral | 1.28 | 2.55 | -3.71 | 6.28 | 0.6147 |
| | cVNS | Ipsilateral | 0.00 | . | | | . |
| | taNS | Contralateral | 0.00 | . | | | . |
| | taNS | Ipsilateral | 0.00 | . | | | . |

**Supplementary Table S6.** LMM beta estimates for LC amplitude

| **Effect** | | | **Beta Estimate** | **Standard Error** | **Lower CL** | **Upper CL** | **Pr > \|t\|** |
|---|---|---|---|---|---|---|---|
| **Intercept** | | | 1.30 | 1.85 | -2.32 | 4.93 | 0.5124 |
| **Stimulation site** | cVNS | | 3.97 | 1.78 | 0.48 | 7.45 | 0.0256 |
| | taNS | | 0.00 | . | | | . |
| **Brain region** | PC | | -0.48 | 0.57 | -1.60 | 0.63 | 0.3971 |
| | PFC | | -1.38 | 0.55 | -2.46 | -0.29 | 0.0127 |
| | SM | | 0.00 | . | | | . |
| **# of pulses** | Intermediate [5-20) | | 0.18 | 0.54 | -0.88 | 1.24 | 0.741 |
| | Long 20+ | | 0.77 | 0.95 | -1.09 | 2.62 | 0.4178 |
| | Short [1-4] | | 0.00 | . | | | . |
| **Intensity** | 2000+ | | 2.74 | 0.98 | 0.83 | 4.65 | 0.005 |
| | [1000-1500) | | 2.20 | 0.94 | 0.35 | 4.05 | 0.02 |

| | | | | | | | |
|---|---|---|---|---|---|---|---|
| | [1500-2000) | | 2.89 | 1.13 | 0.68 | 5.11 | 0.0104 |
| | <1000 | | 0.00 | . | | | . |
| **Pulse width** | 100 | | -1.08 | 0.53 | -2.13 | -0.04 | 0.0421 |
| | 200 | | 0.00 | . | | | . |
| **Frequency** | High 100+ Hz | | 2.04 | 1.02 | 0.04 | 4.05 | 0.0459 |
| | Medium [30-100 Hz] | | 1.50 | 0.90 | -0.26 | 3.26 | 0.0947 |
| | Low [0-30 Hz) | | 0.00 | . | | | . |
| **Laterality** | Contralateral | | 0.05 | 0.46 | -0.84 | 0.95 | 0.9084 |
| | Ipsilateral | | 0.00 | . | | | . |
| **Stimulation site x Brain region** | cVNS | PC | 2.51 | 0.82 | 0.91 | 4.11 | 0.0021 |
| | cVNS | PFC | 4.17 | 0.80 | 2.59 | 5.74 | <.0001 |
| | cVNS | SM | 0.00 | . | | | . |
| | taNS | PC | 0.00 | . | | | . |
| | taNS | PFC | 0.00 | . | | | . |
| | taNS | SM | 0.00 | . | | | . |
| **Stimulation site x # of pulses** | cVNS | Intermediate [5-20) | 1.04 | 0.78 | -0.49 | 2.57 | 0.1843 |
| | cVNS | Long 20+ | -1.04 | 1.33 | -3.65 | 1.57 | 0.4342 |
| | cVNS | Short [1-4] | 0.00 | . | | | . |
| | taNS | Intermediate [5-20) | 0.00 | . | | | . |
| | taNS | Long 20+ | 0.00 | . | | | . |
| | taNS | Short [1-4] | 0.00 | . | | | . |
| **Stimulation site x Intensity** | cVNS | 2000+ | -5.68 | 1.55 | -8.72 | -2.64 | 0.0003 |
| | cVNS | [1000-1500) | -4.94 | 1.18 | -7.25 | -2.64 | <.0001 |
| | cVNS | [1500-2000) | -6.89 | 1.35 | -9.54 | -4.23 | <.0001 |
| | cVNS | <1000 | 0.00 | . | | | . |
| | taNS | 2000+ | 0.00 | . | | | . |
| | taNS | [1000-1500) | 0.00 | . | | | . |
| | taNS | [1500-2000) | 0.00 | . | | | . |
| | taNS | <1000 | 0.00 | . | | | . |
| **Stimulation site x Pulse width** | cVNS | 100 | -0.25 | 0.87 | -1.95 | 1.45 | 0.7724 |
| | cVNS | 200 | 0.00 | . | | | . |
| | taNS | 100 | 0.00 | . | | | . |
| | taNS | 200 | 0.00 | . | | | . |
| **Stimulation site x Frequency** | cVNS | High 100+ Hz | 1.23 | 1.55 | -1.80 | 4.26 | 0.4256 |
| | cVNS | Medium [30-100 Hz] | -1.37 | 1.44 | -4.18 | 1.45 | 0.3417 |
| | cVNS | Low [0-30 Hz) | 0.00 | . | | | . |
| | taNS | High 100+ Hz | 0.00 | . | | | . |
| | taNS | Medium [30-100 Hz] | 0.00 | . | | | . |

| | | | | | | | |
|---|---|---|---|---|---|---|---|
| | taNS | Low [0-30 Hz) | 0.00 | . | | | . |
| **Stimulation site x Laterality** | cVNS | Contralateral | 0.71 | 0.66 | -0.58 | 2.00 | 0.282 |
| | cVNS | Ipsilateral | 0.00 | . | | | . |
| | taNS | Contralateral | 0.00 | . | | | . |
| | taNS | Ipsilateral | 0.00 | . | | | . |

**Supplementary Table S7.** LMM beta estimates for LC latency

| Effect | | | Beta Estimate | Standard Error | Lower CL | Upper CL | Pr > \|t\| |
|---|---|---|---|---|---|---|---|
| **Intercept** | | | 318.93 | 17.49 | 284.65 | 353.21 | <.0001 |
| **Stimulation site** | cVNS | | 25.00 | 21.86 | -17.84 | 67.84 | 0.2528 |
| | taNS | | 0.00 | . | | | . |
| **Brain region** | PC | | -7.17 | 6.64 | -20.19 | 5.85 | 0.2806 |
| | PFC | | -0.78 | 6.30 | -13.12 | 11.57 | 0.9017 |
| | SM | | 0.00 | . | | | . |
| **# of pulses** | Intermediate [5-20) | | -6.77 | 6.28 | -19.07 | 5.54 | 0.2814 |
| | Long 20+ | | -4.21 | 10.25 | -24.31 | 15.88 | 0.6811 |
| | Short [1-4] | | 0.00 | . | | | . |
| **Intensity** | 2000+ | | -24.51 | 12.37 | -48.75 | -0.27 | 0.0477 |
| | [1000-1500) | | -6.85 | 12.30 | -30.95 | 17.26 | 0.5779 |
| | [1500-2000) | | -23.39 | 13.75 | -50.33 | 3.56 | 0.0891 |
| | <1000 | | 0.00 | . | | | . |
| **Pulse width** | 100 | | 28.80 | 6.31 | 16.42 | 41.17 | <.0001 |
| | 200 | | 0.00 | . | | | . |
| **Frequency** | High 100+ Hz | | 17.32 | 13.44 | -9.03 | 43.67 | 0.1979 |
| | Medium [30-100 Hz] | | 21.70 | 12.64 | -3.08 | 46.47 | 0.0862 |
| | Low [0-30 Hz) | | 0.00 | . | | | . |
| **Laterality** | Contralateral | | -11.40 | 5.30 | -21.79 | -1.01 | 0.0316 |
| | Ipsilateral | | 0.00 | . | | | . |
| **Stimulation site x Brain region** | cVNS | PC | 7.80 | 9.21 | -10.26 | 25.85 | 0.3977 |
| | cVNS | PFC | 0.45 | 8.80 | -16.80 | 17.70 | 0.9592 |
| | cVNS | SM | 0.00 | . | | | . |
| | taNS | PC | 0.00 | . | | | . |
| | taNS | PFC | 0.00 | . | | | . |
| | taNS | SM | 0.00 | . | | | . |
| **Stimulation site x # of pulses** | cVNS | Intermediate [5-20) | 18.73 | 8.69 | 1.70 | 35.76 | 0.0313 |
| | cVNS | Long 20+ | 18.02 | 13.70 | -8.84 | 44.88 | 0.1886 |
| | cVNS | Short [1-4] | 0.00 | . | | | . |

| | | | | | | | |
|---|---|---|---|---|---|---|---|
| | taNS | Intermediate [5-20) | 0.00 | . | | | . |
| | taNS | Long 20+ | 0.00 | . | | | . |
| | taNS | Short [1-4] | 0.00 | . | | | . |
| **Stimulation site x Intensity** | cVNS | 2000+ | -19.40 | 15.38 | -49.54 | 10.75 | 0.2074 |
| | cVNS | [1000-1500) | 3.19 | 14.45 | -25.12 | 31.51 | 0.8251 |
| | cVNS | [1500-2000) | -10.64 | 15.85 | -41.72 | 20.43 | 0.5022 |
| | cVNS | <1000 | 0.00 | . | | | . |
| | taNS | 2000+ | 0.00 | . | | | . |
| | taNS | [1000-1500) | 0.00 | . | | | . |
| | taNS | [1500-2000) | 0.00 | . | | | . |
| | taNS | <1000 | 0.00 | . | | | . |
| **Stimulation site x Pulse width** | cVNS | 100 | -21.59 | 9.17 | -39.56 | -3.63 | 0.0186 |
| | cVNS | 200 | 0.00 | . | | | . |
| | taNS | 100 | 0.00 | . | | | . |
| | taNS | 200 | 0.00 | . | | | . |
| **Stimulation site x Frequency** | cVNS | High 100+ Hz | -61.13 | 18.50 | -97.39 | -24.88 | 0.001 |
| | cVNS | Medium [30-100 Hz] | -52.02 | 17.59 | -86.49 | -17.55 | 0.0031 |
| | cVNS | Low [0-30 Hz) | 0.00 | . | | | . |
| | taNS | High 100+ Hz | 0.00 | . | | | . |
| | taNS | Medium [30-100 Hz] | 0.00 | . | | | . |
| | taNS | Low [0-30 Hz) | 0.00 | . | | | . |
| **Stimulation site x Laterality** | cVNS | Contralateral | 22.31 | 7.34 | 7.93 | 36.69 | 0.0024 |
| | cVNS | Ipsilateral | 0.00 | . | | | . |
| | taNS | Contralateral | 0.00 | . | | | . |
| | taNS | Ipsilateral | 0.00 | . | | | . |

**Supplementary Table S8.** Output of the LMM for ta-control vs. taNS, for amplitude and latency of the early component (EC), intermediate component (IC) and late component (LC)

| Outcome | Factor Level | LS-Mean* | 95% Confidence Interval | | p-value | ICC |
|---|---|---|---|---|---|---|
| EC, Latency | taCON | 63.21 | 56.24 | 70.18 | 0.9839 | 0.0141 |
| | taNS | 63.28 | 60.64 | 65.92 | | |
| IC, Latency | taCON | 136.97 | 127.59 | 146.34 | 0.0388 | 0.0315 |
| | taNS | 146.37 | 141.89 | 150.84 | | |
| LC, Latency | taCON | 319.31 | 295.11 | 343.52 | 0.8303 | 0.0073 |
| | taNS | 316.63 | 307.40 | 325.86 | | |
| EC, Amplitude | taCONT | 9.16 | 5.77 | 12.54 | <0.0001 | 0.3846 |
| | taNS | 5.74 | 2.58 | 8.90 | | |
| IC, Amplitude | taCONT | 7.75 | 5.72 | 9.79 | <0.0001 | 0.1185 |

| | | | | | | |
|---|---|---|---|---|---|---|
| | taNS | 4.76 | 3.18 | 6.35 | | |
| LC, Amplitude | taCONT | 5.58 | 2.92 | 8.24 | 0.0475 | 0.2800 |
| | taNS | 4.33 | 1.93 | 6.72 | | |

** Least square means estimates were adjusted for the random effect of monkey and fixed effects of stimulation modality, brain region, number of pulses, intensity, frequency, and laterality, along with two-way interactions between stimulation site and each fixed effect.*
